# Medullary epithelium-free areas in the rat thymus are specialized niches enriched for mature thymocytes and distinct stromal subsets

**DOI:** 10.64898/2026.06.21.733571

**Authors:** Yasushi Sawanobori, Tadayuki Ogawa

## Abstract

The thymic medulla provides the microenvironment for negative selection, late thymocyte maturation, and thymocyte egress, and is generally characterized by widespread distribution of medullary thymic epithelial cells (mTECs). In contrast, rat thymic medulla contains medullary epithelium-free areas (mEFAs), but the cellular composition and functional significance of these regions remain unclear. Here, we combined spatial transcriptomics and scRNA-seq, using robust cell-type decomposition (RCTD) to characterize mEFAs in Lewis-strain rat thymus. These analyses revealed that more mature-phenotypes of CD4SP, CD8SP, and regulatory T-cell-lineage thymocytes were preferentially localized in mEFAs, whereas immature SP subsets were enriched in medullary epithelium-containing areas. Newly found rat thymic mesenchymal cell-3 and -4 (TMC3 and TMC4) subsets were also enriched in mEFAs. These subsets were broadly similar to mouse medullary fibroblasts but displayed distinct predicted interactions with SP thymocytes, including costimulatory molecule– receptor, chemokine–receptor, and ECM–integrin axes. In addition, the venous endothelial cells (vECs) expressing portal endothelial cell markers were accumulated in mEFAs. The S1P transporter gene *Spns2* was preferentially expressed in both TMC4 and vEC subsets, suggesting increased local concentration in mEFAs. These findings indicate that rat mEFAs are specialized medullary niches linking stromal organization, thymocyte maturation, and thymic egress.

## Introduction

The thymus is the primary organ for T cell development. Following positive selection, thymocytes (T cell progenitors) undergo further selection in the thymic medulla, where self-reactive cells are eliminated through negative selection^1,2^. During this process, thymocytes are exposed to tissue-restricted antigens (TRAs) presented by medullary thymic epithelial cells (mTECs) and dendritic cells (DCs). Thymocytes that bind these self-antigens with excessively high affinity are deleted, thereby helping to prevent autoimmune diseases. Consistent with this central role, mTECs are distributed throughout the thymic medulla in humans and mice.

In contrast, we previously reported that rat thymic medulla contains medullary epithelium-free areas (mEFAs)^3,4^. Given the importance of mTECs in negative selection, the presence of such areas without overt pathology is difficult to explain. As a preliminary step toward identifying stromal cell subsets localized in mEFAs, we performed immunohistochemistry (IHC) and found mEFA-localized PDGFRβ expression. Because PDGFRβ is a marker for thymic fibroblasts^5^, this finding led us to consider fibroblasts as candidate stromal cells enriched in mEFAs.

Although mTECs are indispensable for T cell development, other stromal populations, including mesenchymal cells and vascular endothelial cells (ECs), also make major contributions to thymic structure and function^6,7^. Mesenchymal lineage cells, including fibroblasts, adventitial cells, and pericytes, are essential during thymic organogenesis. Neural crest-derived mesenchymal cells direct the transformation of the third pharyngeal pouch and surrounding epithelial cells, thereby shaping the thymic primordium^8^. In addition, transcriptional and humoral factors produced by mesenchymal cells support thymic epithelial integrity and normal thymic development^9–12^. Medullary fibroblasts (mFbs) have also been reported to express TRAs, and impaired maturation of these cells can induce autoimmunity^5,13^.

Mesenchymal stromal subsets also play major roles in thymic egress. Two pathways are particularly well established: the CCL21–CCR7 axis and the sphingosine-1-phosphate (S1P)–S1PR1 axis^6^. Although CCL21 is expressed by mTECs, heparan sulfate on adventitial cells and pericytes surrounding blood vessels captures CCL21 and helps attract mature thymocytes. Disruption of the CCL21–CCR7 axis impairs thymic egress and causes defective negative selection and dacryoadenitis^14,15^.

Regulation of the S1P–S1PR1 axis is complex^16,17^. Sphingosine kinase 1 and 2 (*Sphk1* and *Sphk2*) produce S1P. Sphingosine-1-phosphate transporter (Spns2) transports the produced S1P to extracellular spaces. Sphingosine-1-phosphate lyase 1 (*Sgpl1*) degrades intracellular S1P to regulate S1P production to appropriate levels. Finally, phospholipid phosphatase 3 (*Plpp3*) on the plasma membrane dephosphorylates extracellular S1P to maintain the gradient. Mesenchymal cells regulate the S1P– S1PR1 axis via expression of at least two genes: *Sphk1*^*18*^ and *Sgpl1*^*19*^. These reports suggest that the production of S1P from mesenchymal cells – not at excessive levels – is required.

While mesenchymal cells contribute to thymic egress, endothelial cells play significant roles in both thymic entry and egress. These specialized endothelial cells are termed thymic portal endothelial cells (TPECs). Entry-associated TPECs promote progenitor entry through adhesion molecules such as P-selectin, VCAM-1, and ICAM-1^20,21^. On the other hand, egress-associated TPECs regulate thymic egress via S1P: EC-specific deletion of *Spns2* disrupts thymic egress^22–25^. Interestingly, endothelial expression of *Plpp3* is likewise required for proper thymic egress^26^. This apparent paradox suggests that different endothelial subsets may cooperate to establish the S1P gradient required for efficient egress.

However, entry-associated TPECs and egress-associated TPECs remain difficult to distinguish. In a published scRNA-seq dataset, both appear to fall within the same *Selp*^*+*^*Bst1*^*+*^ cluster, even though BST-1^+^ vessels have been proposed as sites of thymic emigration^27^.

These findings led us to hypothesize that mEFAs are not merely structural defects, but specialized medullary regions with distinct stromal organization and function. To test this idea, we sought to identify the differentiation stages of thymocytes and the stromal subsets localized within mEFAs. However, conventional IHC in rat tissue is constrained by the limited availability of rat-reactive antibodies, and many commercially available antibodies do not perform reliably in rat thymic sections. We therefore applied spatial transcriptomics (ST) to obtain an antibody-independent view of mEFA-associated cell populations. At the time this study was conducted, no ST platform optimized specifically for rat tissue was available, and we therefore used a species-independent, reverse-transcription-based platform. Although this approach is applicable to rat samples, transcript capture per cell-equivalent segment is limited, making it difficult to discriminate closely related cell subsets or cellular states from ST data alone. To address this limitation, we generated reference scRNA-seq datasets and used robust cell-type decomposition (RCTD)^28^ to infer cellular composition within the ST data. By combining this approach with IHC, we identified accumulation of mature thymocyte subsets, uniquely differentiated mesenchymal subsets, and egress-associated TPECs within mEFAs, thereby clarifying the functional characteristics of these unusual medullary regions.

## Results

### Rat thymic overview and ST data acquisition

We used the ED21 antibody to depict medullary areas, as this antibody specifically binds to rat mTECs^4^ and its antigens are presumably keratins 14 and 17 (discussed later). As we previously reported, the thymic medulla of 8–9-week-old Lewis rats contains mTEC-deficient mEFAs containing delicate type IV collagen fibers^3,4^ (Fig. 1A). We further assessed age- and strain-dependent variation in mEFA extent. In Lewis rats, mEFAs were limited at 3 and 5 weeks of age, expanded by 8–9 weeks, and remained comparably extensive at 6 months. At 8–9 weeks, DA and ACI rats exhibited smaller mEFAs than Lewis rats, whereas F344 rats showed a similar extent.

**Fig. 1:**
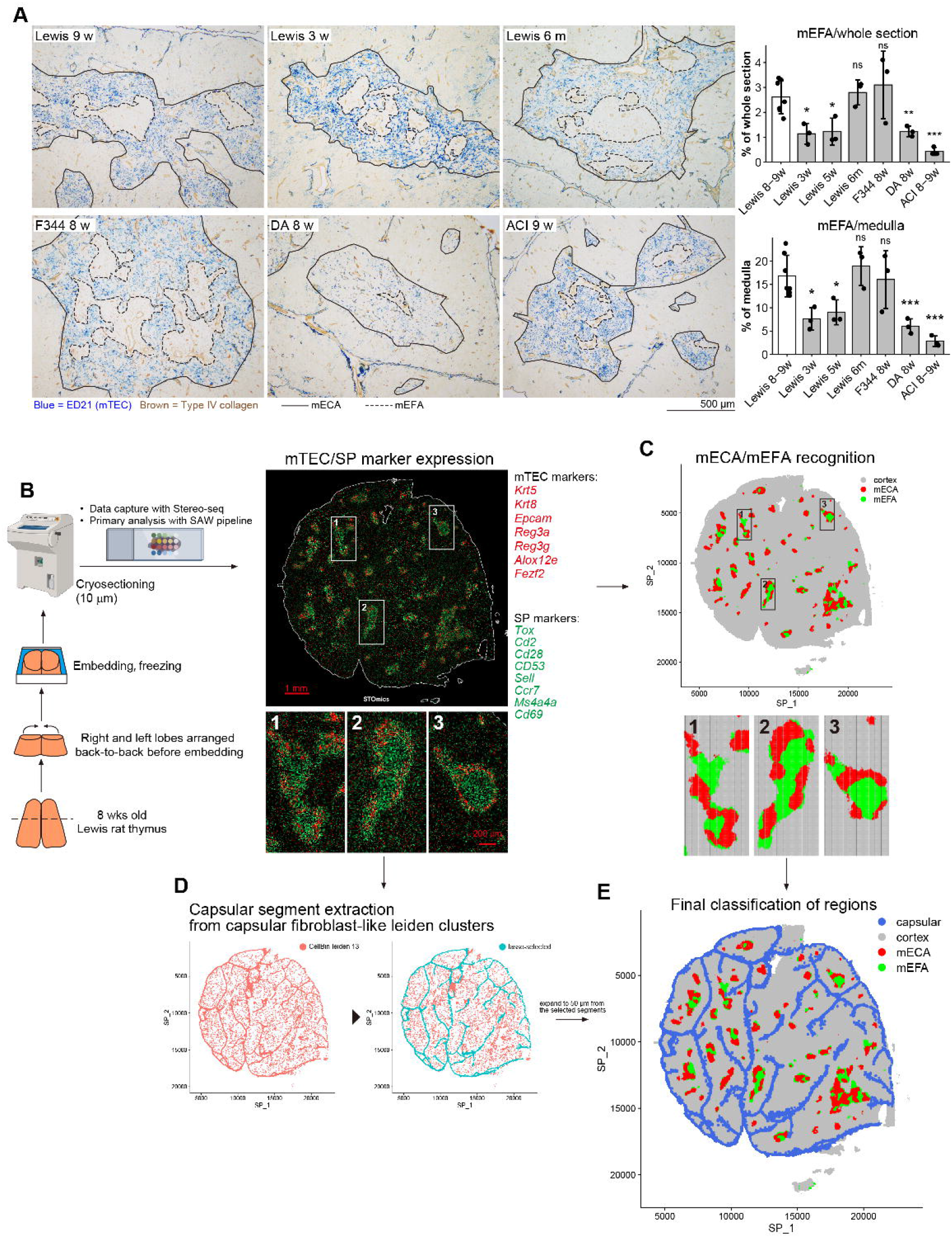
Rat thymic overview and ST data acquisition. A: Representative pictures of ED21-immunohistochemically stained sections from different ages and strains and counted areas of mEFA regions. Statistical significance was examined with Student’s t-test. ns: not significant, *: *p* < 0.05, **: *p* < 0.005, ***: *p* < 0.0005. B: Schematic overview of the workflow for spatial transcriptomics and detected mECAs and medullas. A male 8-week-old Lewis rat was used. mECAs and medullas are depicted as mTEC- and SP thymocyte-associated gene-expressing areas, respectively. Gene expression was visualized using StereoMap. C: mECAs, mEFAs, and cortex regions identified computationally using Seurat in the ST dataset. D: Leiden clusters containing capsular fibroblasts and lasso-selected capsular segments. E: Regions determined in the annotation procedures. C, D, and E depict Bin20_obj.

To spatially define these regions, we performed Stereo-seq on a thymic section from an 8-week-old Lewis rat and processed the data using the Stereo-seq Analysis Workflow pipeline (Fig. 1B). This yielded two h5ad-formatted spatial transcriptomics datasets: an ssDNA-based cell-segmented dataset (CellBin) and a uniformly segmented dataset at 20-bin (10 μm) resolution (Bin20). We annotated medullary compartments based on marker expression, defining SP-marker^hi^ mTEC-marker^hi^ regions as medullary epithelium-containing areas (mECAs) and SP-marker^hi^ mTEC-marker^low^ regions as mEFAs. The combined area of these two regions was considered the medulla.

The CellBin and Bin20 datasets were imported into Seurat, where mTEC and SP marker scores were calculated and smoothed by k-nearest-neighbor averaging to facilitate recognition of mECAs and mEFAs (Fig. 1C). Capsular regions, including both the outer capsule and interlobular capsular regions, were identified using capsular fibroblast-like Leiden clusters (Fig. 1D). Capsular segments were manually selected and then expanded by 50 μm to generate capsular-region annotations. These annotations were then combined to generate the final spatial map used for downstream analyses (Fig. 1E).

### Generation of reference scRNA-seq datasets and cell type identification

As references for cell type identification in the ST dataset, we performed scRNA-seq for rat thymocytes (Fig. S1A) and thymic stroma (Fig. S2A). However, the numbers of cells analyzed were too small to identify clusters for each T-cell developmental pathway or mTEC subtype. Therefore, we further deployed published mouse datasets for projection mapping.

We obtained an scRNA-seq dataset of 8.5 × 10^3^ thymocytes from a Lewis rat (Fig. S1A). The dataset was projected onto UMAP space, and clusters were identified with typical markers for developmental stages. As a mouse thymocyte reference, we used a dataset comprising 7.7 × 10^4^ cells from CD4-fated, CD8-fated, and wild-type animals^29^. We regenerated clusters referring to the marker molecules mentioned in the original article, with several modifications (Fig. S1B). First, we subdivided the CD4 and CD8 conventional T-cell developmental pathways into more detailed subsets based on the distributions of CD4- or CD8-fated thymocytes and the representative transcriptional factors *Zbtb7b* and *Runx3* (Fig. S2C). Second, the *Ikzf2*^+^*Bcl2l11*^+^*Nr4a1*^+^ cluster, designated Neg. Sel. (1) in the original paper, was divided into agonist-selected (AgoSel) 1 and 2, as AgoSel1 expressed markers of agonist-selected Treg precursors, including *Nr4a1, Nr4a3, Trib1, Cd69, Nfkb1*, and *Nfkbia*^30,31^ (Fig. S2D). In contrast, AgoSel2 expressed genes characteristic of gut intra-epithelial lymphocyte precursors, including *Pdcd1, Nrgn*, and *Cd160*^31^. Third, the *Tnfrsf4*^+^*Tnfrsf18*^+^ Neg. Sel. (2) cluster in the original reference was renamed to Pre-Treg based on its proximity to the Treg cluster. The rat and mouse thymocyte datasets were integrated using the JointPCAIntegration algorithm, and the mouse clustering was projected onto the rat cells (Fig. S1E).

For rat thymic stromal scRNA-seq, we magnetically sorted CD45-cells from the thymus of a Lewis rat, performed sequencing, and mapped the cells onto the UMAP space (Fig. S2A). Capsular fibroblasts (capFb), vascular smooth muscle cells and pericytes (vSMC/PC), and endothelial cells (Endo) were identified based on previous rat and mouse literature^6,7,32-34^. TECs were identified with conventional rat TEC markers^3,4,35^. Clusters located between capFb and vSMC/PC were designated thymic mesenchymal cells (TMC). The TMC, Endo, and TEC clusters were temporarily labeled TMC1–4, Endo-1/2, and TEC-a–e, respectively. As a mouse thymic stroma reference, we used a dataset comprising 3.0 × 10^5^ cells from animals of various ages and genotypes with original clustering and UMAP space^36^. The rat and mouse stromal datasets were integrated using the CCAIntegration algorithm, and mouse cluster identities were projected onto the rat cells (Fig. S2B). However, capFb, vSMC/PC, and TMC1–4 clusters were retained with their labels for more rigorous cross-species comparative analyses (Fig. 3). The similarities between the original mouse clusters and the projected rat clusters were confirmed with Pearson correlation (Fig. S2C).

The rat thymocyte and stromal objects with projected clusters were consolidated into a single reference object (Fig. S3A). Some unreliable clusters (intCD4, mature cycling T, NKT, gdT, MEC, nmSC, TEPC, aaTEC1, aaTEC2, mimetic TEC subsets, and fetus-derived cells) were removed due to their scarcity and/or their distance from the original mouse clusters. Based on this refined reference object, cell types of the segments in the CellBin and Bin20 ST datasets were allocated using RCTD.

In both the CellBin and Bin20 datasets, segments predicted as thymocyte lineages were more numerous than those predicted as stromal lineages when only the top-1 RCTD prediction was considered (Fig. S3B). However, thymocytes showed top2/top1 ratios (defined as the number of segments assigned to each cluster as the second RCTD candidate divided by the number assigned as the first candidate) below 1.00, whereas many stromal lineages showed ratios above 1.00. These observations suggested that stromal transcripts were often captured within segments whose major transcriptomic component was thymocytic. Indeed, mTEC-derived genes (including *Cd74*, keratins, and MHC class II genes), TMC-derived genes (including *C7, C4a, S100a5*), and EC-derived genes (Cst3 and Timp3) were enriched in segments predicted as mECA- and mEFA-localized thymocytes, respectively (Fig. S3C; the full gene list is provided in Table S1).

We interpreted this pattern as reflecting unavoidable transcriptomic mixing between thymocytes and adjacent stromal cells. In the CellBin dataset, ssDNA-based segmentation may not fully capture thin TEC or fibroblast protrusions, nor the flattened morphology of endothelial cells, causing transcripts from these cells to be assigned to neighboring thymocyte-dominant segments (Fig. S3D). In the Bin20 dataset, transcriptomic mixing is expected by design because each segment represents a fixed 10-μm spatial bin. However, this uniform segmentation was expected to make stromal transcript contamination less dependent on cell shape and therefore more suitable for comparing the spatial distribution of stromal subsets.

Based on this reasoning, we used the CellBin dataset for thymocyte localization analyses and considered only top-1 RCTD predictions. For analyses of stromal subsets, we used the Bin20 dataset. For clusters with moderate top2/top1 ratios, only top-1 predictions were used. For clusters with particularly high top2/top1 ratios (> 4.2), top-2 predictions were also included because restricting the analysis to top-1 predictions would leave too few segments for stable spatial analysis. These clusters were capFb, TMC1, TMC2, TMC4, early Pr, cTEC, and capEC (Table 1). The resulting distributions of TEC clusters were then visualized, demonstrating the expected cTEC localization in capsular region/cortex and mTEC localization in mECAs (Fig. S3E).

**Table 1:** RCTD predictions in the ST dataset included in analyses.

| dataset | RCTD prediction | Clusters |  |  |
| --- | --- | --- | --- | --- |
|  |  | Thymocyte lineage | TMC3, vSMC/PC, mTEC1, mTEC-prol, mTEC2, aEC, vEC | capFb, TMC1, TMC2, TMC4, early Pr, cTEC, capEC |
| CellBin | Top 1 | analyzed | not analyzed | not analyzed |
|  | Top 2 | not analyzed | not analyzed | not analyzed |
| Bin20 | Top 1 | not analyzed | analyzed | analyzed |
|  | Top 2 | not analyzed | not analyzed | analyzed |

#### Mature SP thymocytes were preferentially localized in mEFAs

The ratios of RCTD-predicted cell types in each area (capsular area, cortex, mECA, and mEFA) were calculated. As is well established, the DP subset was predominantly distributed in capsular areas and the cortex, whereas DP-sig (DP thymocytes that had received the positive-selection signal) appeared to have initiated migration toward the medulla (Fig. 2A).

**Fig. 2:**
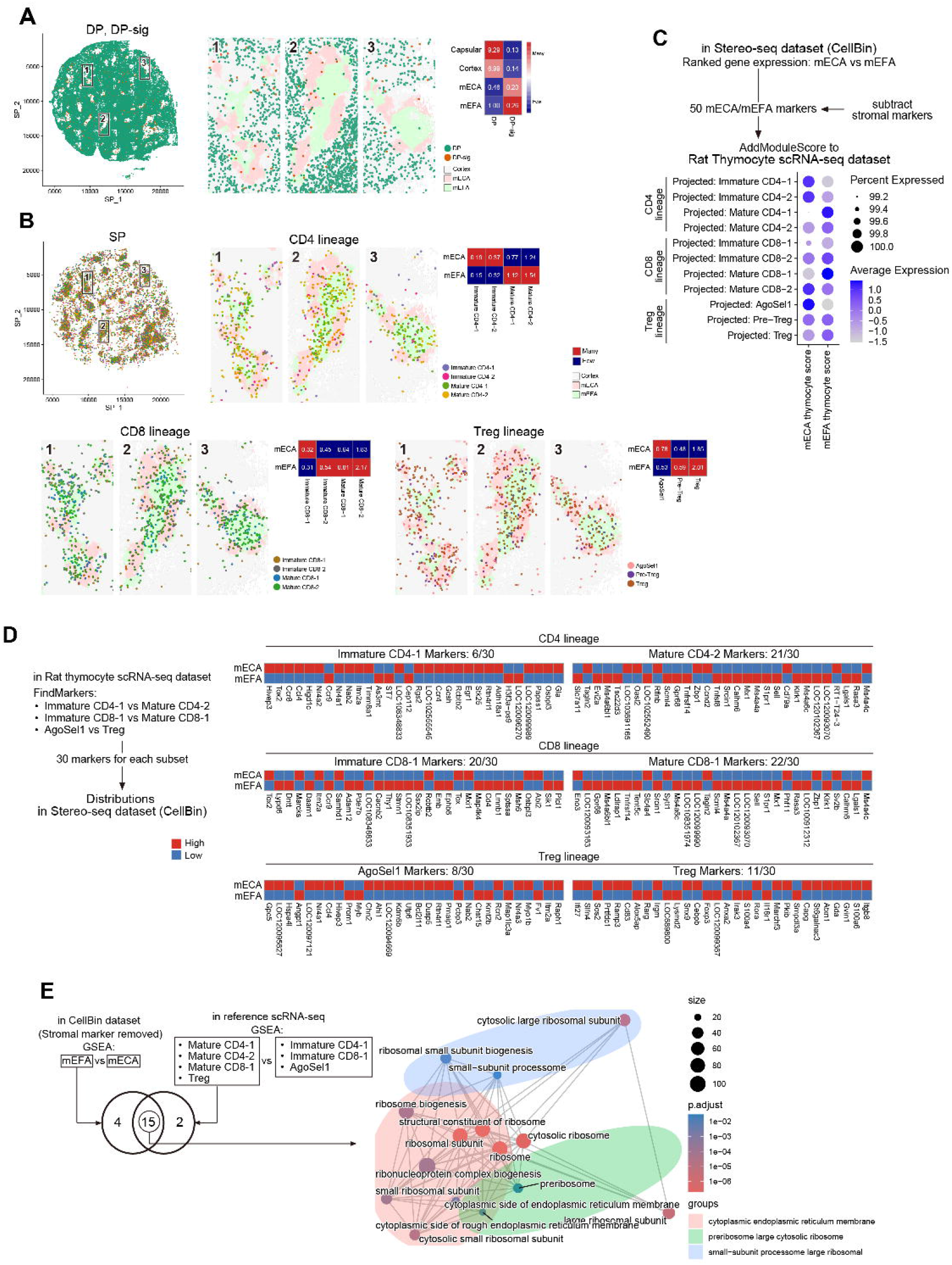
Distribution of thymocyte subsets. A: Distribution of RCTD-predicted segments corresponding to DP and DP-sig subsets in the ST dataset. Digits in the heatmap are % in each region, while colors correspond to z-scores across regions. B: Distribution of RCTD-predicted segments corresponding to SP subsets in the ST dataset. Digits in the heatmap are % in each region, while colors are binarized abundance across mECA and mEFA. C: mECA- and mEFA-thymocyte scores within the scRNA-seq reference dataset, based on markers extracted from the ST dataset. D: Regional expression of immature- and mature-thymocyte markers within the ST dataset. Markers were extracted from the reference dataset. E: Commonality between GSEA results from the ST dataset and the reference scRNA-seq dataset. The commonly detected terms are depicted as an enrichment plot.

Within the SP thymocyte clusters, immature CD4-1, immature CD4-2, immature CD8-1, and AgoSel1 clusters were distributed predominantly in mECA whereas mature CD4-1, mature CD4-2, immature CD8-2, mature CD8-1, mature CD8-2, Pre-Treg, and Treg subsets exhibited mEFA-skewed localization (Fig. 2B). In particular, the enriched Treg distribution in mEFAs corresponded with our previous research ^3^. To further examine the mature SP thymocyte localization in mEFAs, we defined gene sets predominantly expressed in either mEFA or mECA from the ST dataset. These gene sets were then applied to the rat thymocyte scRNA-seq dataset using the AddModuleScore function (Fig. 2C). Consistent with Fig. 2B, the mature CD4-1, mature CD4-2, mature CD8-1, Pre-Treg, and Treg subsets exhibited higher mEFA-thymocyte scores, whereas immature CD4-1, immature CD4-2, immature CD8-1, and AgoSel1 clusters exhibited higher mECA-thymocyte scores. Although the mature CD8-2 cluster is classified as a mature subset, it displayed a high mECA-thymocyte score. Given its lower expression of *S1pr1* and *Sell* (Fig. S1D), the mature CD8-2 cluster was not considered a typical egressing subset; therefore, we designated the mature CD8-1 subset as the representative model of mature CD8SP thymocytes.

Furthermore, we extracted 30 immature or mature markers for each lineage from the rat thymocyte scRNA-seq dataset and surveyed their expression in the ST dataset (Fig. 2D). In all lineages, the numbers of mature markers predominantly expressed in mEFA were greater than the numbers of these predominantly expressed in mECA, reinforcing the mature thymocyte localization in mEFAs.

Finally, to corroborate this “mature thymocytes to mEFA” tendency, we compared two GSEA results: mEFA versus mECA in the ST dataset and mature versus immature SP thymocytes in the rat thymocyte scRNA-seq dataset (Fig. 2E). Fifteen of the 21 significantly enriched GO terms were shared, most of which were ribosome-associated. This concordance is interpreted as independent support for the preferential localization of mature SP thymocytes in mEFAs. The list of all detected GO terms is provided as Table S2.

Of note, because RCTD-predicted thymocyte segments included stroma-derived genes (Fig. S3C), we subtracted these genes from the region-enriched gene sets (Fig. 2C) and the GSEA (Fig. 2E).

#### Rat TMC3/4 subsets were localized in mEFAs, corresponding to mouse medullary fibroblasts, but with unique marker expression patterns

To examine how rat and mouse thymic mesenchymal clusters correspond to each other, the CCA-integrated mesenchymal subsets were compared (Fig. 3A). Consistent with the Pearson correlation analysis (Fig. S2C), TMC1/2 and TMC3/4 clusters were considered to be counterparts of mouse intermediate fibroblasts (intFb) and medullary fibroblasts (mFb), respectively. However, a fraction of cells in the rat TMC4 cluster did not overlap with the mouse mFb cluster, suggesting that TMC4 has a partially distinct gene-expression pattern. The mouse aging-associated TEC2 (aaTEC2) cluster, which exhibits an intermediate gene expression profile between fibroblast and TEC clusters ^36^ (Fig. S3C), was located near, but did not overlap with, the rat TMC4 cluster.

**Fig. 3:**
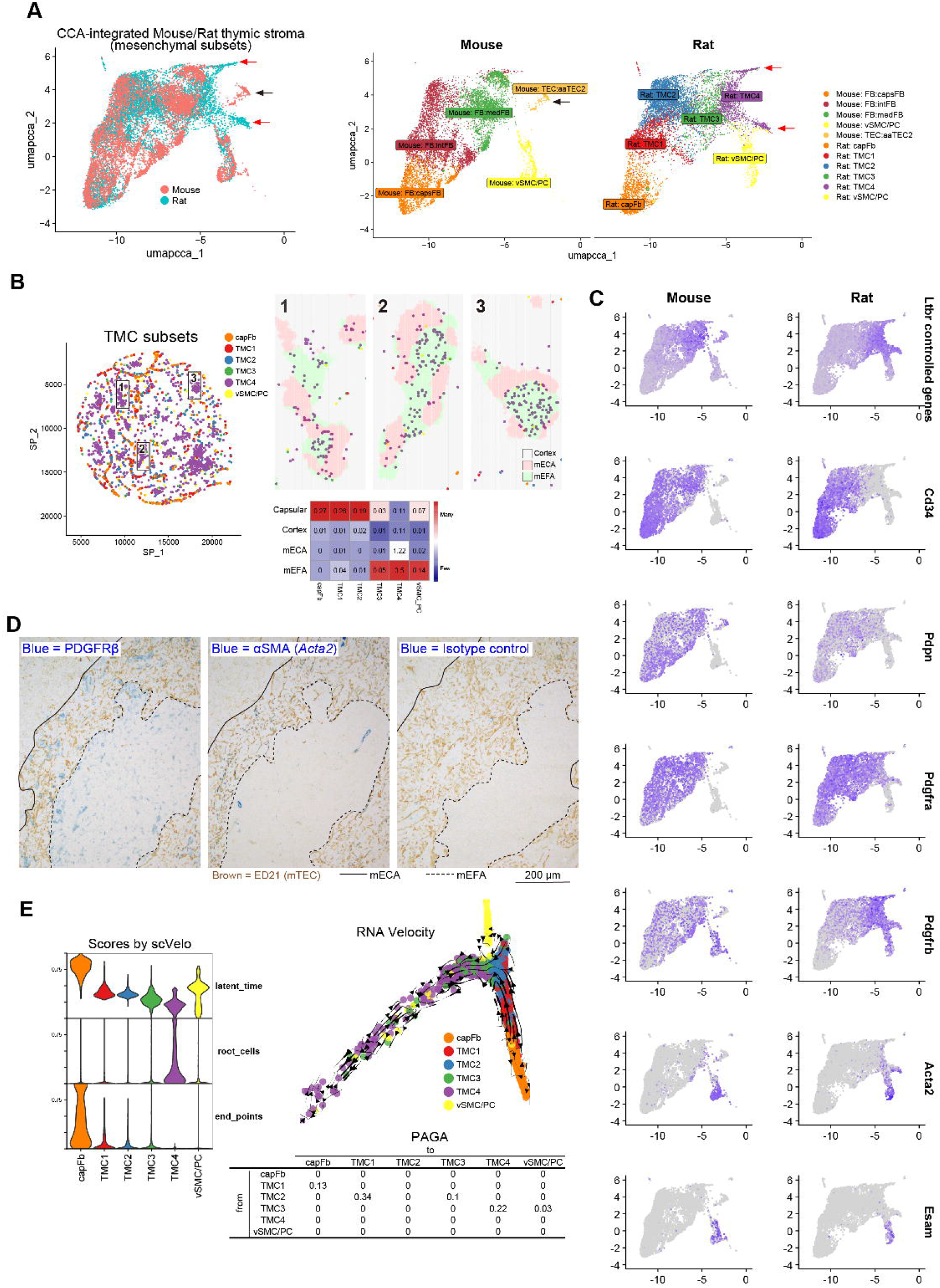
Characterization of mesenchymal subsets and their distribution in the ST dataset. A: Mouse and rat mesenchymal subsets within the interspecifically integrated scRNA-seq datasets of thymic stroma. The integration and UMAP projection are based on the CCAIntegration algorithm. Black and red arrows indicate mouse- and rat-unique cells, respectively. B: Distribution of RCTD-predicted segments corresponding to mesenchymal subsets in the ST dataset. Digits in the heatmap are % in each region, while colors correspond to z-scores across regions. C: Expression of representative marker genes in mouse and rat mesenchymal subsets. The *Ltbr*-controlled genes are provided in Table S3. D: Pictures of anti-PDGFRβ- or anti-αSMA-immunohistochemically stained sections. A thymus of a male 8-week-old Lewis rat was used. E: RNA velocity analysis of rat mesenchymal subsets. Violin plots of latent time, root cell probability, end-point probability, a UMAP plot of RNA velocity, and a table of PAGA scores are presented.

Next, we investigated the localization of rat mesenchymal subsets in the Stereo-seq dataset (Fig. 3B). capFb and TMC1/2 clusters were mainly localized in capsular regions, whereas TMC3/4 and vSMC/PC clusters were enriched in mEFAs. Lymphotoxin β receptor-dependent genes in mFb are thought to function as mFb-derived TRAs^5,7,13^; they were expressed in TMC3/4 (Fig. 3C; the list of genes is provided in Table S3). At the same time, several fibroblast marker molecules showed species-specific expression patterns. Although *Cd34* and *Pdgfrb* were expressed in both intFb and mFb clusters in mice, *Cd34* expression was absent from rat TMC3/4, whereas *Pdgfrb* expression was largely restricted to TMC3/4. *Acta2* and *Esam* were specifically expressed in vSMC/PC in both mouse and rat. Based on these marker specificities, we performed immunohistochemistry to further confirm the localization of the TMC clusters (Fig. 3D). PDGFRβ protein was detected mainly in mEFAs, consistent with the localization of TMC3 and TMC4. αSMA (*Acta2*) staining was observed around blood vessels within mEFAs, corresponding to vSMC/PC. αSMA expression was also detected in a subset of mTECs, as suggested in the initial analysis (Fig. S2A).

Finally, to investigate the transcriptional states of TMC clusters, we performed RNA velocity analysis (Fig. 3E). The TMC4 cluster exhibited the highest root cell probability, while the capFb cluster exhibited the highest latent time and end-point probability. Velocity streamlines projected onto the UMAP showed bidirectional flows between TMC4 and TMC2/3, indicating heterogeneous transcriptional dynamics among these subsets. Partition-based graph abstraction (PAGA) analysis suggested two transition pathways: TMC2 → TMC1 → capFb and TMC3 → TMC4. Taken together, these analyses suggest that capFb represents a terminal remodeling state, whereas TMC4 occupies an upstream transcriptional position. However, the velocity signals among TMC2, TMC3, and TMC4 were mixed, preventing a clear linear ordering of these subsets.

### A cell–cell interaction analysis and S1P-associated gene expression revealed distinct roles of rat TMC3/4 subsets in T cell development and egress

To predict the roles of rat TMC subsets in T cell development, we performed cell–cell interaction analysis using CellChat^37^. Interactions between all medullary stromal subsets and SP thymocyte subsets were surveyed and displayed (Fig. 4A). The detected interactions were grouped into the categories Chemokine & Receptors, MHC – CD4/CD8, Costimulatory Signals, Cytokines & Receptors, Notch Signaling, Adhesion & Retention, ECM & Basement Membrane, ECM-binding growth factors, and Synapse-like Adhesion. We identified a substantial number of interactions that were unique to rat TMC3/4–SP thymocyte pairs. Notably, the ECM & Basement Membrane category contained the largest number of these interactions. A complete list of detected interactions is provided in Table S4. The rat-specific interactions were neither seen in the mouse result before ortholog conversion, indicating these differences were not caused by the conversion procedure.

**Fig. 4:**
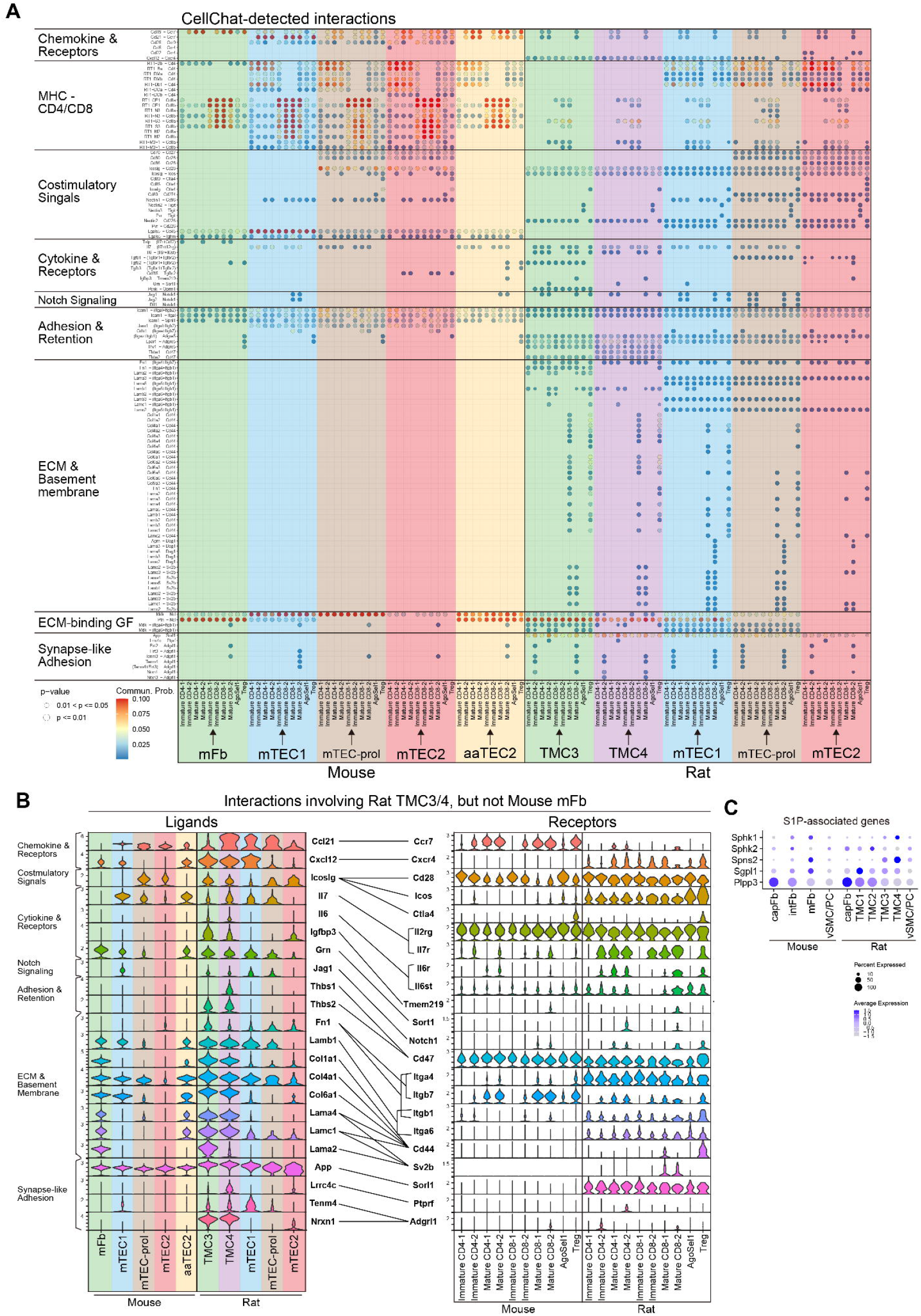
CellChat-detected interactions between thymic stromal subsets and SP thymocytes, and S1P-associated gene expression. A: CellChat-detected interactions between thymic stroma and SP thymocytes. Results from mouse and rat datasets are combined and displayed in a single table. B: Violin plots of ligand and receptor genes of interactions involving rat TMC3/4, but not mouse mFb subsets. C: Expression of S1P-associated genes in mouse and rat mesenchymal subsets. Expression is scaled in each species. Note that mouse gene symbols are converted to rat orthologs in A and B.

Next, interactions involving rat TMC3/4 and SP subsets, but not the mouse mFb subset, were extracted. Ligand and receptor genes underlying these interactions were then examined by violin plots (Fig. 4B). Some ligand genes, such as *Ccl21, Icoslg, Il7*, and *Thbs1/2*, were expressed in rat TMC3/4 but absent from mouse mFbs. In other cases, ligands were expressed in both rat TMC3/4 and mouse mFbs, whereas the corresponding receptors, such as *Cxcr4, Sort1, Itga6, Cd44*, Sv2b, and *Sorl1*, were expressed in rat SP subsets but absent from mouse SP subsets, suggesting that both rat stromal and SP subsets have distinct developmental programs from mouse counterparts. Notably, among these receptors, *Itga6, Cd44*, and *Sv2b*, which bind to laminins and collagens, accounted for the large number of rat-specific stroma-SP interactions in the ECM & Basement Membrane category.

Moreover, some ligands, including *Il6, Thbs1/2*, collagens, laminins, *Lrrc4c*, and *Nrxn1*, were not shared with mTEC lineage subsets. This suggests that TMCs not only compensate for mTECs’ functions but also have their own unique functions. Interestingly, *Ccl21* and *Il7* were expressed not only in rat TMC3/4 but also in the mouse aaTEC2 subset. This observation raises the possibility that some functions of rat TMC3/4 may be performed by aaTEC2 cells in mice.

As stated in the Introduction, the S1P–S1PR1 axis is essential for thymic egress. Therefore, we also examined S1P-associated gene expression in mesenchymal clusters (Fig. 4C). In both rat and mouse, *Sphk1* and *Spns2* were preferentially expressed in TMC4/mFb clusters, whereas *Plpp3* was expressed in capFb clusters. These differences may contribute to spatial compartmentalization of S1P-metabolic programs. Interestingly, *Sgpl1* was preferentially expressed in TMC1 cluster in rat and mFb cluster in mouse, suggesting that a different S1P gradient is required in the rat thymus, possibly due to extensive mEFAs.

### mEFAs contained thymic portal endothelial cells associated with thymic egress

In the current consensus, murine ECs are largely divided into capsular EC (capEC), venous EC (vEC), and arterial EC (aEC)^36,38^. Our thymic stromal scRNA-seq dataset also included these three phenotypes (Fig. 5A). The lymphatic EC marker *Lyve1* was expressed in only a small number of cells and was therefore considered negligible (data not shown).

**Fig. 5:**
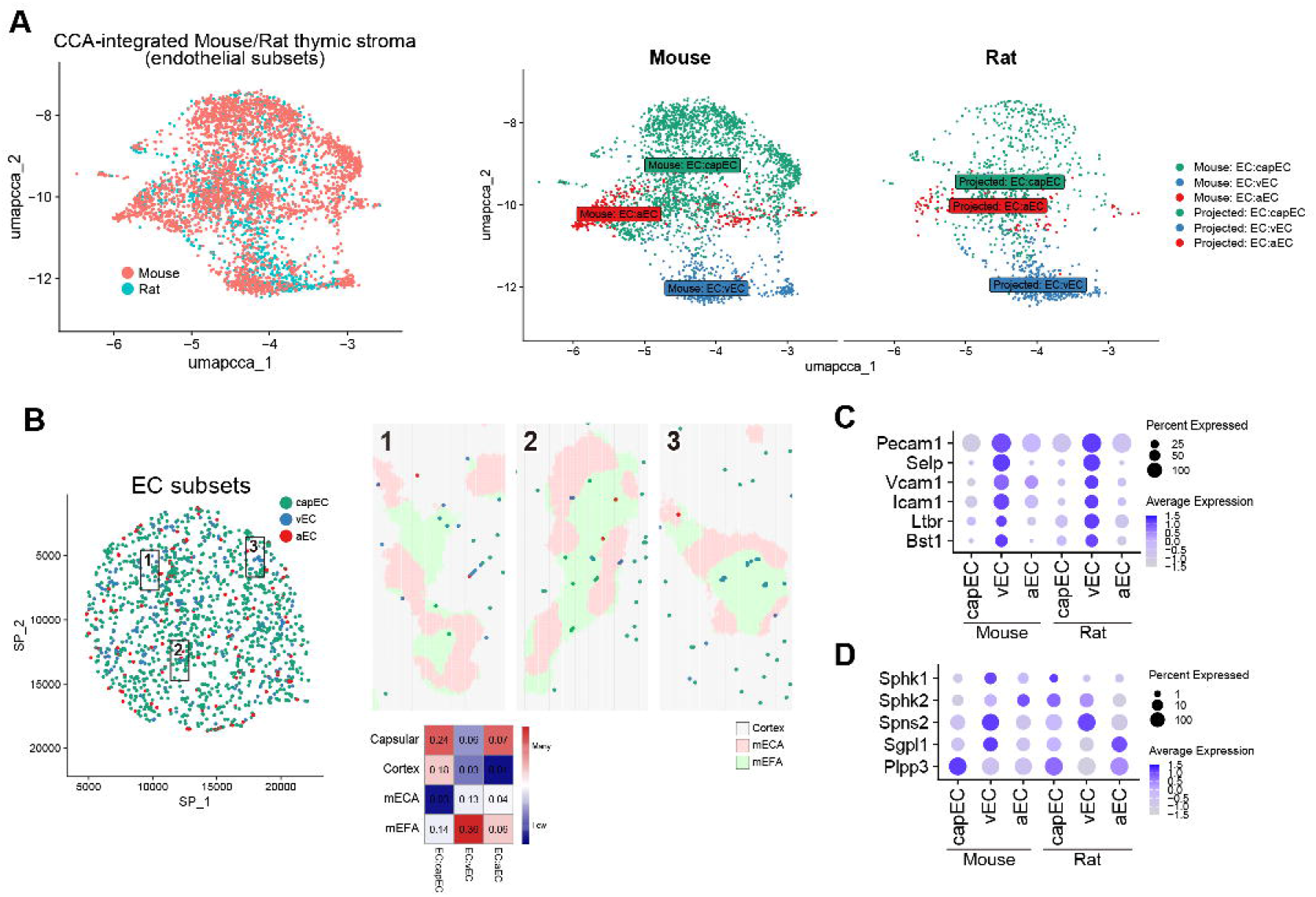
Characterization of endothelial subsets and their distribution in the ST dataset. A: Mouse and rat endothelial subsets within the interspecifically integrated scRNA-seq datasets of thymic stroma. The integration and UMAP projection are based on the CCAIntegration algorithm. B: Distribution of RCTD-predicted segments corresponding to endothelial subsets in the ST dataset. Digits in the heatmap are % in each region, while colors correspond to z-scores across regions. C: Expression of TPEC-associated genes in mouse and rat endothelial subsets. D: Expression of S1P-associated genes in mouse and rat endothelial subsets. In C and D, expression is scaled in each species.

RCTD analysis of the ST dataset revealed that capECs were mainly distributed in capsular areas, whereas vECs and aECs were distributed predominantly in mEFAs (Fig. 5B).

TPECs express P-selectin (*Selp*), ICAM-1, and VCAM-1, and require signaling via LTβR to exert full potency^6,20,21,39^. Therefore, we investigated expression of these transcripts within EC clusters. All of these genes were expressed in the vEC cluster (Fig. 5C). Although TPECs have been reported as entry sites for T cell progenitors, it is also known that BST-1^+^ TPECs function as exit sites for mature thymocytes^27^. Consistent with this, *Bst1* was also preferentially expressed in the vEC cluster. However, it was difficult to distinguish entry- and egress-associated TPECs in the scRNA-seq dataset.

Because endothelial cells also regulate the S1P–S1PR1 axis, we examined S1P-associated genes in EC clusters (Fig. 5D). *Spns2* was preferentially expressed in vECs, whereas *Plpp3* was predominantly expressed in capECs. Thus, S1P export- and degradation-associated genes were segregated between distinct EC subsets, with the export-associated program concentrated in vECs, which were abundant in the mEFAs.

### Identification of candidate ED21 target proteins

In our previous study, we showed that ED21 preferentially stains MHCII^low^AIRE-mTECs, and that the apparent molecular weight of the ED21-reactive protein is approximately 50 kDa^4^ (Fig. S4A). In the present scRNA-seq reference dataset, MHC class II- and *Aire*-related genes were preferentially expressed in the mTEC2 cluster, suggesting that the ED21-reactive antigen is more strongly associated with mTEC1-like cells than with mTEC2 cells.

To identify candidate ED21 target proteins, rat thymic lysates were immunoprecipitated with biotin-conjugated ED18, ED19, ED21, OX3, or isotype-control IgM antibodies, followed by DIA-MS analysis of the precipitated samples (Fig. S4B). The complete list of detected protein groups is provided in Table S5. As a positive control, OX3 immunoprecipitation resulted in the detection of RT1-Ba and RT1-Bb protein groups, confirming that our procedure could recover known antibody targets. Candidate ED21 antigens were screened as ED21-specific, molecular weight-matched protein groups, and examined in the reference scRNA-seq dataset (Fig. S4C). Among the candidates, *Krt14, Idh1, Krt42*, and *Krt17* showed TEC-associated expression. Of these, *Krt14* and *Krt17* were less abundant in the mTEC2 cluster, consistent with the ED21 staining pattern. The spatial expression of these genes was further examined in the ST dataset (Fig. S4D). *Idh1* and *Krt42* did not reproduce the ED21-like spatial pattern, whereas *Krt14* and *Krt17* were concentrated in mECAs. In addition, *Krt14* was detected along capsular regions, corresponding to ED21 staining observed along the capsule by IHC (Fig. S4E), in concordance with the TEC:early Pr cluster (Fig. S3E) and human epithelial stem cells^40^.

Because ED21 also stains mouse thymic sections, we next surveyed the expression of *Krt14, Krt17*, and related paralogs in the rat and mouse reference datasets (Fig. S4F). *Krt14* and *Krt17* were expressed in both mouse and rat TEC clusters. Although *Krt13* and *Krt15* were also expressed in rat TECs, they were not detected among ED21-enriched immunoprecipitated proteins (Table S5), making them less likely candidates under the present experimental conditions.

These findings suggest that ED21 may recognize a keratin-family epitope represented by keratin 14 and keratin 17 in rat and mouse thymus. Sequence alignment revealed highly conserved regions among these proteins, providing a possible explanation for this cross-reactivity (Fig. S4G).

## Discussion

In this study, combining scRNA-seq and spatial transcriptomics with IHC, we showed that rat thymic mEFAs contain accumulated mature SP thymocyte subsets, mesenchymal subsets similar to mouse medullary fibroblasts but with distinct gene-expression profiles, and thymic portal endothelial cells associated with thymic egress.

A fundamental question is why mEFAs develop in rat thymi. In mice, both IL-4Rα-deficient and CD1d-deficient models exhibit mEFA-like structures together with reduced thymic egress, suggesting that interactions between iNKT cell-derived type 2 cytokines and IL-4 receptors on mTECs are required for proper medullary organization and function^41^. In addition, iNKT cells promote the emergence of an AIRE^+^ TEC subset in reaggregated thymic organ cultures^42^, whereas impaired mTEC development reduces the number and activity of iNKT cells^42,43^.

Our findings raise the possibility that a similar relationship operates in rat. Lewis and F344 rats reportedly contain very few thymic iNKT cells, although F344 rats retain detectable iNKT cells in the liver^44^. This is consistent with our observation that both Lewis and F344 rats exhibited significantly expanded mEFAs compared with DA and ACI rats (Fig. 1A). The relative scarcity of TEC-lineage cells compared with mesenchymal-lineage cells in the Lewis thymic stromal dataset may also fit this interpretation (Fig. S2). Notably, the Lewis strain is highly susceptible to a broad range of autoimmune disease models, including adjuvant arthritis^45^, EAE^46^, uveitis^47^, and EAN^48^. One possible explanation for this susceptibility is that iNKT-cell scarcity in the thymus impairs mTEC development and thereby weakens negative selection.

An important issue, however, is whether mEFAs simply represent dysfunctional medullary regions. Our data suggest a different interpretation. First, we observed TMC3/4-specific LTβR-regulated TRA expression (Fig. 3C). Nitta *et al*. reported that mFbs express TRAs and that impaired maturation of these cells can lead to autoimmunity^5^. In our previous study, we also noted that, although reduced in number, DCs were still present in mEFAs at non-negligible levels^3^. Therefore, it is possible that TRAs expressed by TMC subsets are presented to SP thymocytes through DCs within mEFAs. Second, both rat mesenchymal subsets and SP thymocytes appear to acquire specialized features that may preserve medullary function despite the paucity of mTECs in mEFAs (Fig. 4A-B). Interactions between rat TMC3/4 subsets and SP thymocytes were more diverse than those between mouse medullary fibroblasts and SP thymocytes.

Among these interactions in mEFAs, CCL21 and CXCL12 are known to contribute to stage-specific thymocyte positioning^49^. Although CXCL12 has generally been associated with cortical thymocytes because its receptor CXCR4 is preferentially expressed in DN and DP subsets, our analysis indicated that rat SP subsets uniquely express *Cxcr4*. This finding suggests that rat SP thymocytes may have adapted to respond to mesenchymal cues within mEFAs.

The same tendency was observed for ECM-related interactions. ECM components such as fibronectin, laminin, and collagen are well established as important regulators of T-cell development^50–53^, yet CellChat did not detect comparable interactions between mouse medullary stromal cells and SP thymocytes. In contrast, the increased expression of *Itga4, Itga6*, and *Itgb1* in rat SP subsets, together with selective *Cd44* expression in Tregs, produced abundant predicted interactions between TMC3/4 and SP subsets. This pattern suggests that mesenchyme-thymocyte interactions are particularly important in the mEFA-containing rat medulla.

TMC3/4 subsets also expressed *Icoslg* and *Il7*, ligands shared with mTEC subsets and known to be required for thymocyte differentiation, negative selection, and survival in the medulla^54–57^. This finding raises the possibility that rat TMC3/4 subsets partially compensate for the local absence of mTECs in mEFAs by acquiring mTEC-like supportive functions.

At the same time, rat TMC3/4 subsets expressed ligands not shared with either mTECs or mouse medullary fibroblasts, including *Il6, Thbs1/2, Lrrc4c*, and *Nrxn1*. In peripheral tissues, stromal cell-derived IL-6 regulates the balance between Treg and T helper subsets and contributes to tissue remodeling^58,59^. THBS1/2 is deposited into the ECM, where it suppresses T-cell activation and regulates matrix assembly^60–63^. LRRC4C and NRXN1 are best known as synapse-associated adhesion molecules that mediate selective and stable cell-cell contacts^64,65^. Although their roles in the thymus remain unknown, these molecules may help maintain balance among SP subsets within mEFAs, restrain excessive activation, support survival and stable ECM architecture, and reinforce contacts between SP thymocytes and the stromal scaffold.

We observed age-dependent expansion of mEFAs (Fig. 1A): they were small at 3–5 weeks, reached a maximum by 8 weeks, and remained large at least until 6 months. This time course indicates that mEFA expansion is age related, while it begins too early to be regarded simply as senescence. This finding is notable in relation to TMC4, an mEFA-localized subset that showed several similarities to mouse aaTEC2, including proximity on the integrated UMAP (Fig. 3A) and shared expression of *Ccl21* and *Il7* (Fig. 4B). In contrast, TMC4 did not directly overlap with the mouse mFb cluster, making it difficult to regard TMC4 as a straightforward rat counterpart of mouse mFbs. Combining these results with the fact that mouse aaTEC2 was originally identified as epithelial by lineage tracing^36^, it is possible that TMC4 may represent a corresponding population. However, confirmation will require comparable lineage-tracing studies in rats.

As stated in the Introduction, it has been reported that both mesenchymal (mFb and pericytes) and EC subsets contribute to S1P gradient to promote egress of mature thymocytes^18,19,22-26^. Consistent with this concept, we found both TMC4 and vEC expressed the S1P transporter gene *Spns2*, while both capFb and capEC express S1P dephosphorylate gene *Plpp3* (Fig. 4C and 5D). As TMC4 and vEC are accumulated in mEFAs, these results suggest S1P-mediated thymic egress occurs in mEFAs, while S1P concentrations are kept low in capsular regions.

It is highly possible that mEFAs contain exit-associated TPECs (Fig. 5), whereas in mice, exit-associated blood vessels have been reported mainly at the corticomedullary junction (CMJ)^18,27^. In our material, mEFAs did not always coincide with the CMJ (Fig. 1D); some appeared to be surrounded by mTECs. A similar pattern was reported in IL-4Rα-deficient mice, in which mEFA-like structures were also not restricted to the CMJ^41^. In that study, increased labeling by intravenously injected anti-CD4 antibody in IL-4Rα-deficient thymi suggested accumulation of portal endothelial structures and mature thymocytes within mEFA-like regions, consistent with our findings. Together, these observations suggest that mEFAs containing TPECs can form not only near the CMJ but also deep within the medulla.

Given the broad genomic and cellular conservation among mammals, it is unlikely that mEFAs, their associated mesenchymal subsets, and their supporting gene programs are entirely unique to rats. Rather, mice and humans may also have the capacity to generate similar structures under altered signaling conditions, disease states, or senescence. Further investigation of mEFAs and their localized stromal niches may therefore improve our understanding of medullary thymic organization, thymic egress, and the mechanisms linking stromal architecture to immune tolerance and autoimmunity.

## Materials and methods

### Animals

Inbred Lewis, F344, DA, and ACI rats, aged 3–8 weeks, were purchased from SLC Co. (Shizuoka, Japan) and maintained under specific pathogen-free conditions. Some animals were kept in the facility to obtain samples from 6-month-old rats. The exact ages and sexes of the animals used in each experiment are indicated in the corresponding figures, figure legends, and source data. All animal procedures were approved by the Dokkyo Medical University Animal Experiment Committee (approval no. 1358), in accordance with the university’s regulations for animal experiments and Japanese Governmental Law (No. 105).

### Antibodies

Antibodies used for IHC and flow cytometry are listed in Table S6. Some antibodies were purified from culture supernatants and conjugated in house.

### Immunohistochemistry

Tissues were embedded in Tissue-Tek O.C.T. compound (Sakura Finetek Japan, Tokyo, Japan), snap-frozen, and stored at −80°C. Frozen blocks were sectioned at a thickness of 4 μm. After air drying for 2 h and fixation in acetone, sections were stained as described previously^3,4^. Briefly, acetone-fixed sections were rehydrated in Tris-buffered saline (pH 7.4), and then refixed in 4% paraformaldehyde containing 1% calcium chloride solution. After blocking with Block Ace (KAC, Kyoto, Japan), sections were incubated with primary antibodies for 1 h or overnight. After washing with PBS, sections were further incubated with the appropriate secondary antibodies or streptavidin, and signals were developed using Vector Blue substrate (Vector Laboratories, Newark, CA, USA), the New Fuchsin Substrate System (Agilent, Santa Clara, CA, USA), or 3,3’-diaminobenzidine (DAB) substrate (Dojindo Molecular Technologies, Kumamoto, Japan). For multicolor staining, these steps were repeated sequentially.

Photomicrographs were acquired using a BX53 microscope equipped with UPlanFL N objective lenses and a DP27 digital camera (Olympus, Tokyo, Japan). Images were captured with cellSens software (Olympus). Exposure and contrast settings were kept constant within each experiment. For area measurements, montage images of whole thymic sections were generated, and measurements were performed using the cellSens software.

For cross-strain and cross-age analyses, areas containing ED21^+^ thick reticular cells (mTECs) were defined as mECAs, whereas areas lacking mTECs and filled with delicate type IV collagen fibers were defined as mEFAs^3,4^.

### Spatial transcriptomics

A thymus was cut horizontally into three pieces, and the middle portion was embedded in O.C.T. compound in a 1 × 1 cm plastic tray. To maximize the tissue area within the tray, right and left lobes were positioned back to back. A 10-μm-thick section was mounted onto a Chip T (STOmics, Shenzhen, Guangdong, China). The slide was incubated at 37°C for 5 min and then stored in an airtight container at −80°C. Library preparation and sequencing were outsourced to GENEWIZ (Tokyo, Japan). For cell segmentation, ssDNA staining was performed. The resulting FASTQ files and the image of the ssDNA-stained section were processed using the Stereo-seq Analysis Workflow pipeline (ver. 8.1.3, STOmics), with the mRatBN7.2 genome assembly as a reference. For brief visualization of gene expression, StereoMap (ver. 4.1.2, STOmics) was used.

The output files, cellbin_1.0.adjusted.h5ad and bin20_1.0.h5ad, were imported into the R environment (ver. 4.5.1) using the reticulate (ver. 1.43.0), anndata (ver. 0.8.0), and zellkonverter (ver. 1.18.0) packages. Gene expression counts, spatial coordinates, and metadata were extracted and reconstructed as Seurat (ver. 5.3.0) objects, designated CellBin_obj and bin20_obj, respectively. Feature IDs were remapped to gene symbols. Count matrices and coordinates were stored in the obj@assays$RNA@counts and obj@reductions$spatial slots, respectively. The objects were then processed to generate log-normalized expression values (obj@assays$RNA@data). To reduce the object size, z-scored data (obj@assays$RNA@scale.data) were removed.

To identify mECA and mEFA regions, mTEC marker genes and SP thymocyte marker genes (Fig. 1B) were applied to the ST datasets using the AddModuleScore() function. These scores were z-scaled and then smoothed by k-nearest-neighbor averaging using the FNN package (ver. 1.1.4.1). In the CellBin_obj, segments with top 10% of the smoothed SP-thymocyte score and the top 15% of the smoothed mTEC score were defined as mECAs, whereas segments within the top 10% of the smoothed SP-thymocyte score but outside the top 10% of the smoothed mTEC score were defined as mEFAs. In the bin20_obj, the smoothed mTEC score was adjusted to the top 10% based on anatomical concordance with the mECA extent identified in CellBin_obj.

To define capsular regions, capsular segments were manually identified within *Dpp4* and *Pi16*-enriched Leiden clusters, using a custom Shiny (ver. 1.11.1)- and plotly (ver. 4.11.0)-based tool. The selected capsular segments were then spatially dilated by 50 μm. Thus, the resulting capsular annotation corresponds to an approximate total capsular width of 100 μm plus the width of the originally selected segments.

### Tissue digestion

For scRNA-seq analysis of thymocytes, thymi were minced and incubated at 37°C for 25 min in an enzyme mixture containing 0.2% collagenase D (Roche Diagnostics, Indianapolis, IN, USA) and 0.01% DNase I (Roche) in Hank’s buffered saline solution (HBSS) supplemented with 1.25 mM Ca^2+^, 1.45% bovine serum albumin (BSA), and 2% fetal calf serum (FCS). During incubation, the tissues were mechanically dissociated every 5 min by pipetting sequentially with truncated 1000-μL pipet tips, Pasteur pipettes, and heat-tapered tip Pasteur pipettes. After incubation, EDTA was added to a final concentration of 2.5 mM, and the samples were incubated for an additional 5 min. The digested tissue was then filtered through pieces of 50-μm nylon mesh. The resulting cell suspension was centrifuged and resuspended in ACK lysis buffer (145 mM NH_4_Cl, 10 mM KHCO_3_, and 0.1 mM EDTA, pH 7.4). After incubation at 4°C for 10 min, cells were resuspended in PBS, counted using a hemocytometer, and used for each experiment.

For preparation of thymic stromal cells, thymi were digested using a similar procedure, except that the enzyme mixture contained 0.0125% Liberase TH (Roche) and 0.004% DNase I. After the ACK treatment, cells were resuspended at 1.25 × 10^9^ cells/ml in PBS(−) containing 0.5% BSA and 2 mM EDTA (MACS buffer). To block nonspecific binding, ChromPure® mouse IgG whole molecule (Jackson ImmunoResearch, West Grove, PA, USA) was added at 10 μg/ml. After incubation at 4°C for 10 min, anti-rat CD45 MicroBeads (Miltenyi Biotec, North Rhine-Westphalia, Germany) were added at 10% of the suspension volume. After incubation at 4°C for 15 min, the cell suspension was diluted sixfold with MACS buffer then separated using the Depl05 program on an autoMACS instrument (Miltenyi Biotec). The CD45− fraction was stained with 7-AAD (IMMUNOSTEP, Salamanca, Spain), and Pacific Blue-conjugated anti-CD45 antibody (BioLegend, San Diego, CA, USA). Viable CD45− cells were further sorted using a FACSAria III cell sorter (Becton, Dickinson and Company, Franklin Lakes, NJ, USA) and collected in HBSS supplemented with 1.25 mM Ca^2+^, 1.45% BSA, and 2% FCS. The sorted cells were washed three times with PBS(−) and then subjected to scRNA-seq analysis.

### scRNA-seq and reference construction for RCTD

Prepared thymocyte and stromal cell suspensions were submitted to the on-site NGS library construction and sequencing service at GENEWIZ. The resulting FASTQ files were processed using the Cell Ranger pipeline (ver. 8.0.0; 10x Genomics, Pleasanton, CA, USA), with the mRatBN7.2 genome assembly as the reference.

The output filtered_feature_bc_matrix folders were imported into the R environment and converted into Seurat objects, designated RatThymocyte.obj and RatThymicStroma.obj. These objects were then processed according to the standard Seurat workflow^66^.

A published mouse thymocyte dataset was obtained from the Gene Expression Omnibus (GSE186078)29. Expression matrices from individual animals were imported into R and integrated into a multilayer Seurat object using the IntegrateLayers() function with method = JointPCAIntegration. Clustering was performed independently of the original study with some modifications (see Results and Fig. S1B). This object was designated all_m_thymocyte.

A published mouse thymic stroma dataset, ThymoSight, was obtained from Zenodo (12516405)^36^. The file thymosight_cd45neg_TOTAL_mouse.h5ad was imported into R using the zellkonverter package and converted into a Seurat object. The original metadata, clustering, and UMAP coordinates were retained. This object was designated MouseThymicStroma.obj.

To project mouse-derived annotations onto rat cells, gene symbols in the mouse objects were converted to rat orthologs using convert_orthologs() in the orthogene package (ver. 1.14.1), with non121_strategy = “keep_popular” to resolve non-one-to-one mappings without expanding the feature dimension of the expression matrices. Because the function refers to the periodically updated g:Profiler database, all conversions were performed in September 2025, and the resulting ortholog conversion table was saved for reproducibility (Table S7). The symbol-converted all_m_thymocyte and MouseThymicStroma.obj were then integrated with RatThymocyte.obj and RatThymicStroma.obj, respectively, using the IntegrateLayers() function. CCAIntegration and JointPCAIntegration were used respectively.

Mouse cluster labels were then projected onto rat cells using a k-nearest-neighbor-based label transfer procedure in the UMAP space of the integrated Seurat objects. The 50 nearest mouse neighbors of each rat cell were identified using the FNN package (ver. 1.1.4.1). Rat-cell annotation was assigned by majority vote among the neighboring mouse cells. A projected label was accepted only when the most frequent donor label accounted for at least 55% of the neighbors; otherwise, the cell was left unlabeled. The projected labels were then transferred back to RatThymocyte.obj and RatThymicStroma.obj, which were subsequently integrated to generate RatThymicRef. After generating RatThymicRef, several minor populations including NKT, gdT, aaTEC subsets, and mimetic TEC subsets were excluded to reduce the computational cost of RCTD, and because their representation in the rat dataset was too limited to support reliable annotation. The resulting reduced reference object was named RatThymicRef_l and was used for subsequent RCTD analyses.

The similarities between original and projected clusters were calculated as Pearson correlation coefficients. The gene expression matrix was extracted from the interspecifically integrated scRNA-seq dataset of thymic stroma. Standard deviations were calculated among the projected clusters, and genes were ranked by these deviations. Pearson correlation coefficients were calculated with the expression values of top 1000 genes between the original mouse clusters and the projected rat clusters.

### RCTD

RCTD was performed using the spacexr package, according to the standard protocol^67^. The reference object was generated from RatThymicRef_l and the query objects were generated from CellBin_obj and bin20_obj. The run.RCTD() function was executed with doublet_mode = “multi”, allowing assignment of multiple candidate cell types to a single segment. From the resulting RCTD objects, the $cell_type_list, $conf_list, and $sub_weights slots were extracted and added to the metadata of the ST objects. Evaluation of the resulting annotations is described in the Results (Fig. S3).

### Gene set enrichment analysis (GSEA)

To perform GSEA on thymocyte-derived transcriptomes across regions, region-grouped expression profiles were extracted from CellBin_obj using the AverageExpression() function. Genes were ranked according to the expression difference between mEFA and mECA. To reduce contamination from stromal transcripts, marker genes for mTECs, TMC3/4, and EC clusters were identified in RatThymicRef object using FindAllMarkers() and removed from the ranked gene list. GSEA was then performed using clusterProfiler (ver. 4.16.0).

For mature-versus-immature GSEA, the mature and immature SP subsets were combined within RatThymicRef object. Grouped expression data were extracted using AverageExpression(), and genes were ranked according to the expression difference between the mature and immature groups. GSEA was then performed using gseGO() function on the resulting ranked list. As per the default setting, GO terms with an adjusted *p*-value of less than 0.05 were considered statistically significant.

Pathways identified in each GSEA result were compared using a Venn diagram.

### Lymphotoxin β receptor-dependent genes

A bulk RNA-seq dataset including *Ltbr*-deficient mFb samples was downloaded from GSE147357^5^. Gene counts were averaged within each group and transformed to log2 scale. Genes whose average expression in the *Ltbr*-deficient mFb group was at least 2 log2 units lower than that in the control mFb group were selected. These genes were then filtered against a list of tissue-restrictedly expressed genes^68^. The resulting genes were converted to rat orthologs and considered lymphotoxin β receptor-dependent TRAs in mFbs (Table S3). This gene set was then applied to RatMouseStroma.obj using the AddModuleScore() function.

### RNA velocity analysis

RNA velocity analysis was performed on rat mesenchymal subsets according to a standard protocol^69^. A loom file containing spliced and unspliced counts was generated from the rat stromal scRNA-seq output, using velocyto (ver. 0.17.17). The loom file and expression matrix were imported into a Python environment (ver. 3.6.13). After normalization and log transformation, first- and second-order moments were calculated using the neighborhood graph derived from the UMAP embedding. RNA velocity was then estimated using the dynamical model, and velocity vectors were projected onto the UMAP embedding using scVelo (ver. 0.2.4).

To characterize transcriptional dynamics, latent time, root-cell probability, end-point probability, and partition-based graph abstraction (PAGA) were calculated. These velocity-derived metadata were exported and integrated back into the Seurat object for downstream visualization and analysis.

### Cell-cell interaction analysis

Cell-cell interaction analyses were performed using the CellChat package (ver. 2.2.0). For the rat analysis, gene symbols in the mouse interaction database (CellChatDB.mouse) were converted using the orthogene package. Interactions between stromal subsets and mature SP subsets were extracted from the rat and mouse CellChat result objects using the subsetCommunication() function and then compiled into a combined table. Mouse genes in the table were converted to rat orthologs.

### Immunoprecipitation and mass spectrometry

A thymus from an 8-week-old Lewis rat was homogenized in 5 mL of ice-cold lysis buffer containing 50 mM Tris-HCl (pH 7.4), 1% NP-40, 125 mM NaCl, 5 mM EDTA, and cOmplete protease inhibitor cocktail (Roche Diagnostics, Indianapolis, IN, USA), using a manual homogenizer (Fig. S4B). The lysate was incubated on ice for 15 min and centrifuged at 1,000 × g for 10 min. The supernatant was further centrifuged at 10,000 × g for 10 min. The resulting supernatant was adjusted to 10 mL and stored at −80°C until use.

Dynabeads™ M-280 Streptavidin (Invitrogen, Carlsbad, CA, USA) were dispensed into five 5-mL tubes, at 100 μL per tube. After adding 1 mL of PBS, the beads were collected using a DynaMag-5 magnetic rack, and the supernatant was removed. This washing process was repeated once, and the beads were resuspended in 100 μL of 0.1% BSA-PBS. Biotin-conjugated antibodies (ED18, ED19, ED21, or isotype-control IgM; 40 μg each, or OX3, which recognizes rat MHC class II molecules RT1-Ba/Bb^u^, as a positive control for the workflow; 10 μg) were added to the tubes. The suspensions were incubated for 30 min at room temperature with gentle agitation to form bead-antibody complexes. The complexes were washed four times with 0.1% BSA-PBS. Thymic lysate (2 mL) was added to each tube then incubated for 30 min at 4°C with gentle agitation. The bead-antibody-antigen complexes were washed three times and resuspended in 100 μL of elution buffer containing 4 M urea and 10 mM dithiothreitol. After incubation at room temperature for 15 min, the beads were collected using the magnetic rack, and the supernatants were recovered. The elution step was repeated once, and the combined eluates were stored at −80°C until further processing.

Before trypsin digestion, the eluted samples were reduced and alkylated with 30 mM iodoacetamide in 100 mM ammonium bicarbonate. The samples were then adjusted to final concentrations of 1 M urea and 2.5 mM dithiothreitol and digested with sequencing grade modified trypsin (Promega, Madison, WI, USA) for 16 h at 37°C. The resulting peptide were desalted and filtered using C18-Tip devices (CPE C-TIP 1000-C18-S2-100, Nikkyo Technos, Tokyo, Japan) before LC-MS/MS analysis. The processed peptides were analyzed by SWATH acquisition on a TripleTOF 6600 system (SCIEX, Marlborough, MA, USA). The acquired data were then processed using DIA-NN (ver. 2.2.0 Academia)^70^. A rat protein sequence database was prepared from the UniProtKB *Rattus norvegicus* canonical FASTA file. DIA-NN was run in library-free mode. A predicted spectral library was generated from the FASTA file and used for the subsequent DIA search. The following precursor generation settings were used: protease, Trypsin/P; missed cleavages, 1; peptide length, 7–30 amino acids; precursor charge range, 1–4; precursor m/z range, 300–1800; and fragment ion m/z range, 200– 1800. N-terminal methionine excision was enabled. Cysteine carbamidomethylation was set as a fixed modification. Variable modifications were not included in the first-pass analysis. The output was filtered at 1% FDR. Match-between-runs was not used for the final quantitative comparison.

Molecular weights of proteins were automatically obtained from Uniprot and added to the DIA-NN output (Table S5) using an R script (add_uniprot_mw.R). In the DIA-NN output, the antibody-specific enrichment was calculated as:

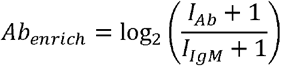

where *I* denotes the DIA-NN-derived protein-group quantity in each immunoprecipitation. The ED21-specificity score was defined as:

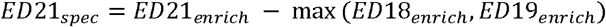

Proteins satisfying the following criteria were considered candidate ED21 antigens and subjected to further investigation: *ED21*_enrich_ > 0, *ED21*_spec_ > 0.3, and 45 < MW (kDa) < 55 (Fig. S4C).

## Supporting information

Original-sized Figures, Supplementary Figures, and Graphical Abstract

## Figure legends

**Fig. S1: Clustering and annotation of the rat thymocyte scRNA-seq dataset**.

A: A UMAP plot and a dot plot of representative markers of the rat thymocyte scRNA-seq dataset. Clustering was performed briefly as an initial analysis.

B: A UMAP plot of a mouse thymocyte scRNA-seq dataset. Expression matrices from multiple individuals were integrated with JointPCAIntegration algorithm. Clustering was performed following the original study with some modifications.

C: Distributions of cells from CD4-fated and CD8-fated transgenic mice, and expression of representative marker genes for CD4^+^ and CD8^+^ SP subsets.

D: Expression of marker genes in the mouse thymocyte scRNA-seq dataset.

E: UMAP plots of the interspecifically integrated scRNA-seq datasets of thymocytes. Datasets were integrated with JointPCAIntegration algorithm, and mouse clustering was projected to rat cells.

**Fig. S2: Clustering and annotation of the rat scRNA-seq dataset of thymic stroma**.

A: A UMAP plot and a dot plot of representative markers of the rat scRNA-seq dataset of thymic stroma. Clustering was performed briefly as an initial analysis.

B: UMAP plots of the interspecifically integrated scRNA-seq datasets of thymic stroma. Datasets were integrated with CCAIntegration algorithm, and mouse clustering was projected to rat cells.

C: Pearson correlation coefficients represent similarities between mouse and rat thymic stromal subsets.

**Fig. S3: RCTD-based prediction of cell subsets**.

A: A UMAP plot of the integrated scRNA-seq dataset of rat thymic stroma-thymocytes. The datasets were integrated with JointPCAIntegration algorithm.

B: The number of predicted segments in the CellBin and Bin20 ST datasets. Top 1 to 3 predictions are displayed. Note that Y-axes are logarithmic.

C: TEC/TMC specific genes detected in RCTD-predicted thymocytes. Segments predicted as thymocytes in mECAs and mEFAs were compared using the FindMarkers() function. Representative enriched genes and their expression in the reference scRNA-seq dataset are displayed.

D: Pictorial representations of ssDNA staining-based segmentation and equal segmentation. Arrows indicate protrusions of fibroblasts and TECs that are contaminated into other segments.

E: Distribution of TEC subsets within the ST dataset. Digits in the heatmap are % in each region, while colors correspond to z-scores across regions.

**Fig. S4: Identification of candidate ED21 target proteins**.

A: ED21 staining pattern reported in our previous study, and MHCII- and *Aire*-related genes in TEC clusters.

B: Schematic overview of the workflow used to narrow down candidate ED21 target proteins.

C: Expression of LC-MS/MS-screened candidate genes in the rat scRNA-seq reference dataset.

D: Spatial expression of candidate genes that showed TEC-associated expression in C, visualized with StereoMap. Arrows indicate *Krt14* expression in capsular regions.

E: Immunohistochemical staining of capsular regions in a male 5-week-old rat thymus. Arrows indicate ED21 staining along an interlobular capsule.

F: Expression of *Krt14* orthologs and paralogs in the mouse and rat scRNA-seq datasets.

G: Protein sequence alignment of mouse and rat keratin 14 and 17.

## Acknowledgement

We thank the members of our laboratory for maintaining the research environment, including reagent management and equipment maintenance. We also acknowledge the use of ChatGPT and Codex for assistance with coding, troubleshooting, and English-language editing. Some illustrations are created with BioRender.com.

## Author contributions

Y.S. conceived and designed the study, performed immunohistochemistry, cell isolation, immunoprecipitation, and computational analyses, curated and interpreted the data, and wrote the manuscript. T.O. performed LC-MS/MS analysis and contributed to study design.

## Data availability

Data supporting this study have been deposited in the Gene Expression Omnibus under accession number GSE333693, including rat thymic stromal and thymocyte scRNA-seq datasets and a spatial transcriptomics dataset from a rat thymic frozen section. The deposited files include scRNA-seq expression matrices, Seurat object metadata, the .loom file used for RNA velocity analysis, h5ad files for spatial transcriptomics, and the .stereo file for StereoMap. The mass spectrometry proteomics data and results have been deposited to the jPOST repository^71,72^ with the dataset identifier JPST004670. R and Python scripts are available on GitHub at https://github.com/yasuhisawanobori/rat_mEFA_spatial_analysis.

This study also used published scRNA-seq datasets for mouse thymocytes (GSE240020) and mouse thymic stroma (https://zenodo.org/records/12516405).

