## Supplementary figures and images for "Medullary epithelium-free areas in the rat thymus are specialized niches enriched for mature thymocytes and distinct stromal subsets"

### Fig1.png

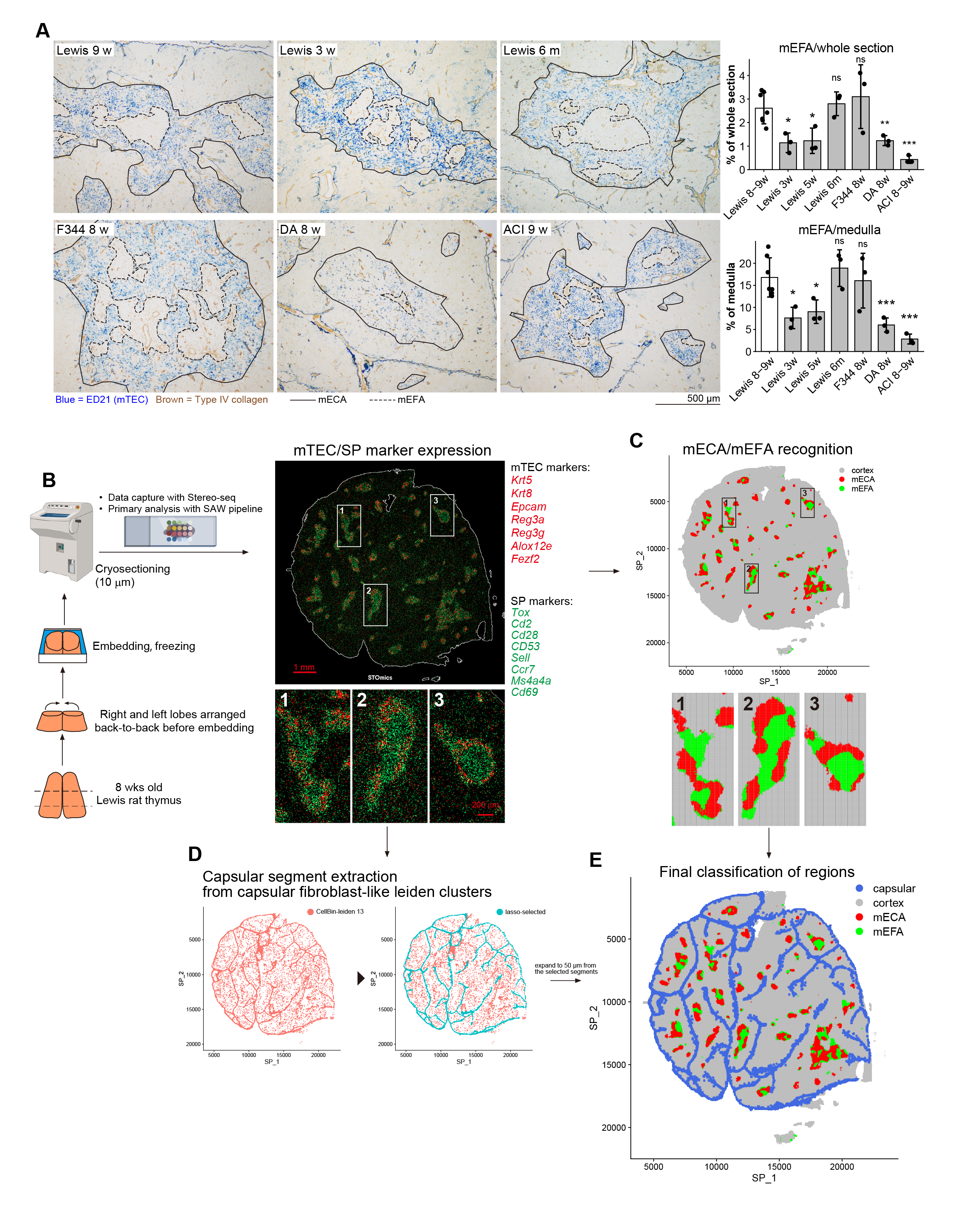

### Fig2.png

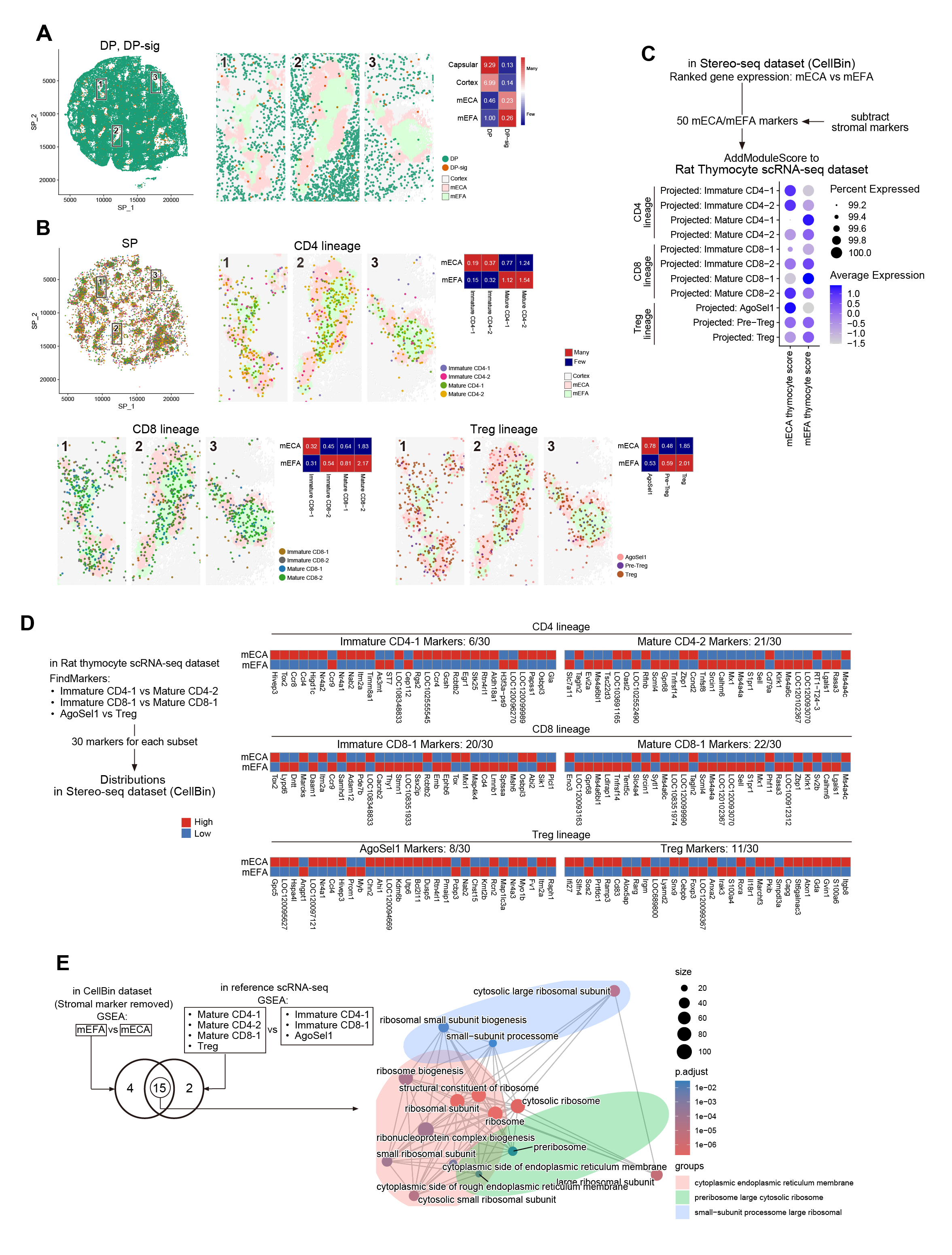

### Fig3.png

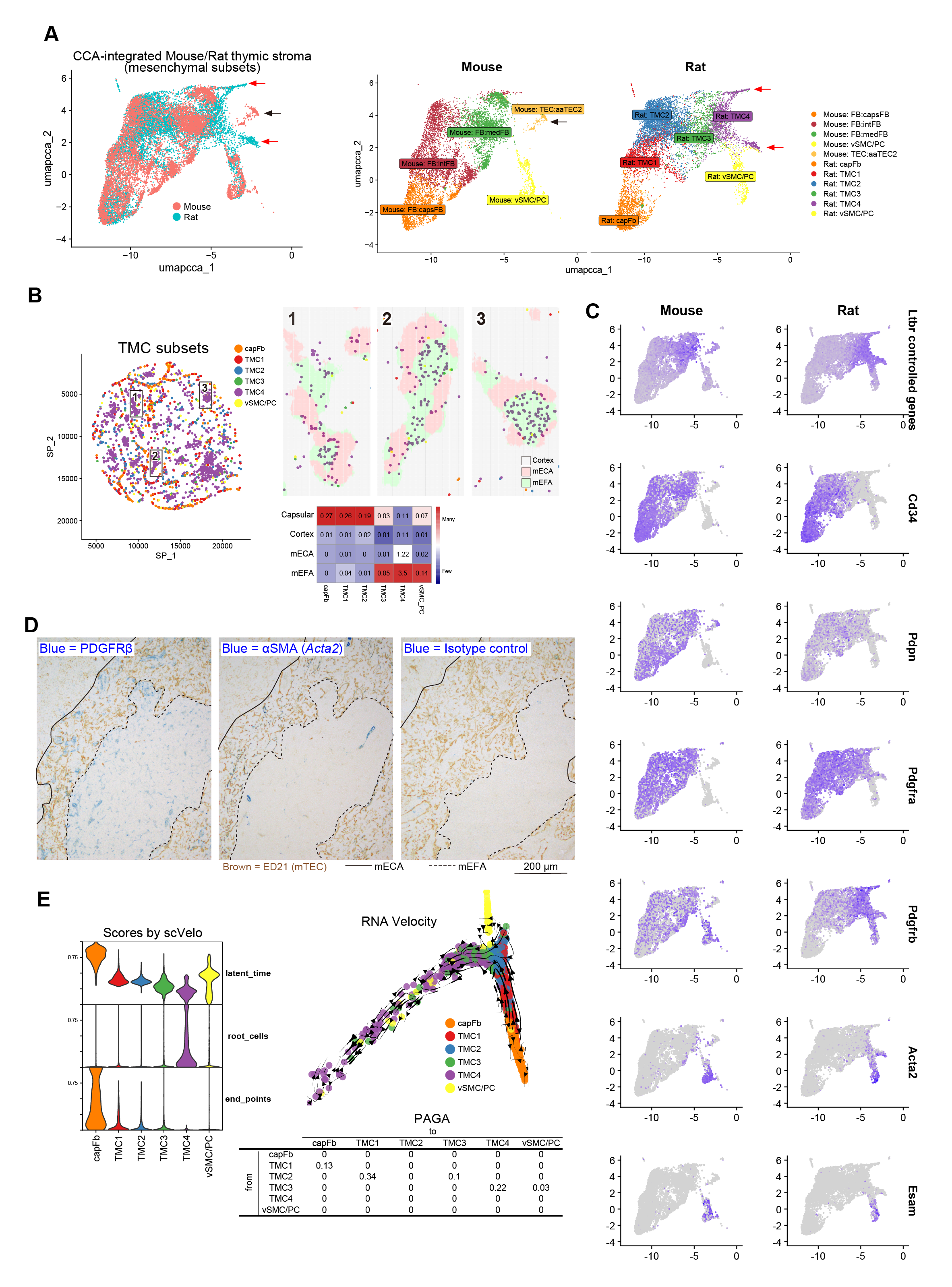

### Fig4.png

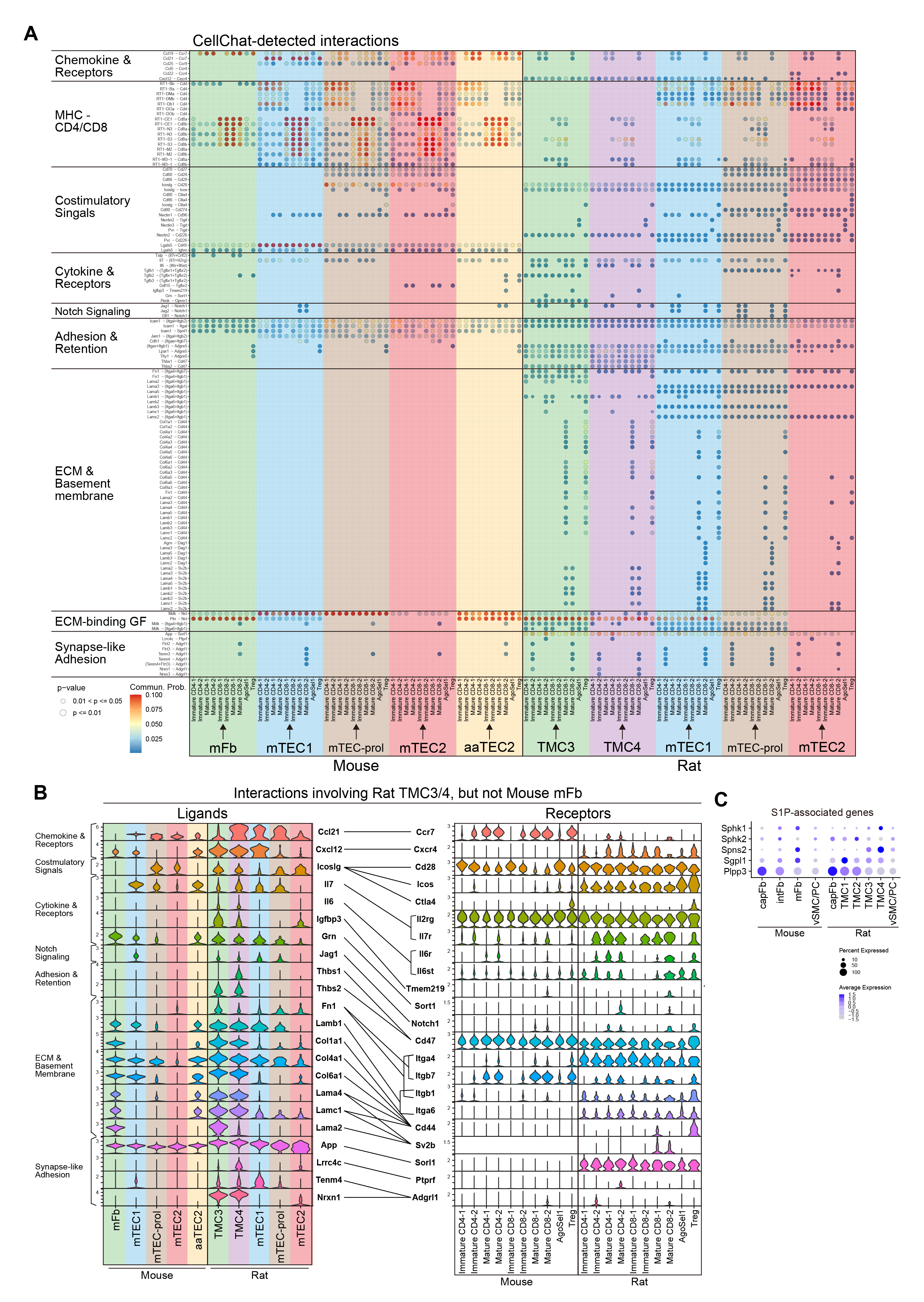

### Fig5.png

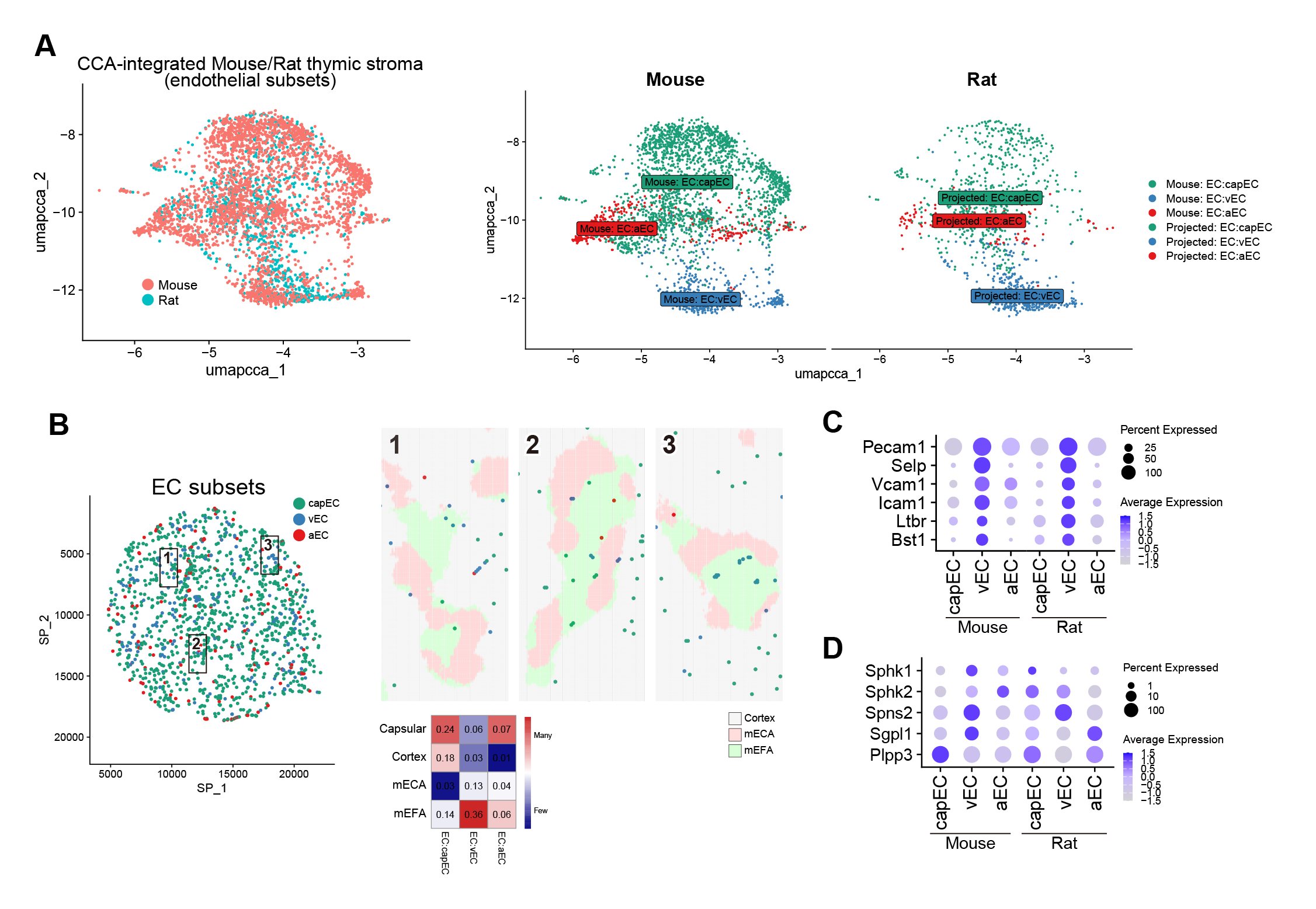

### FigS1.png

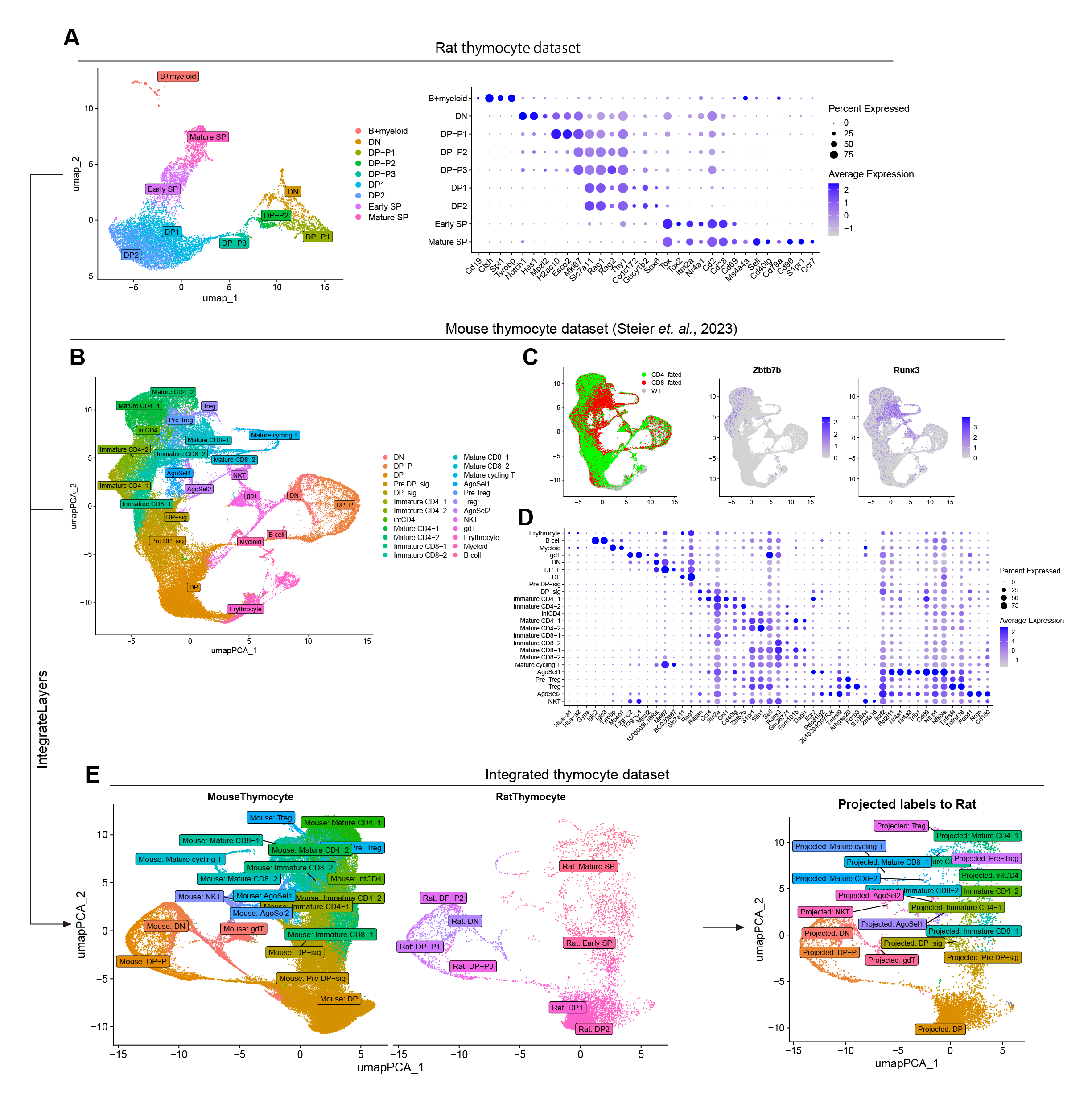

### FigS2.png

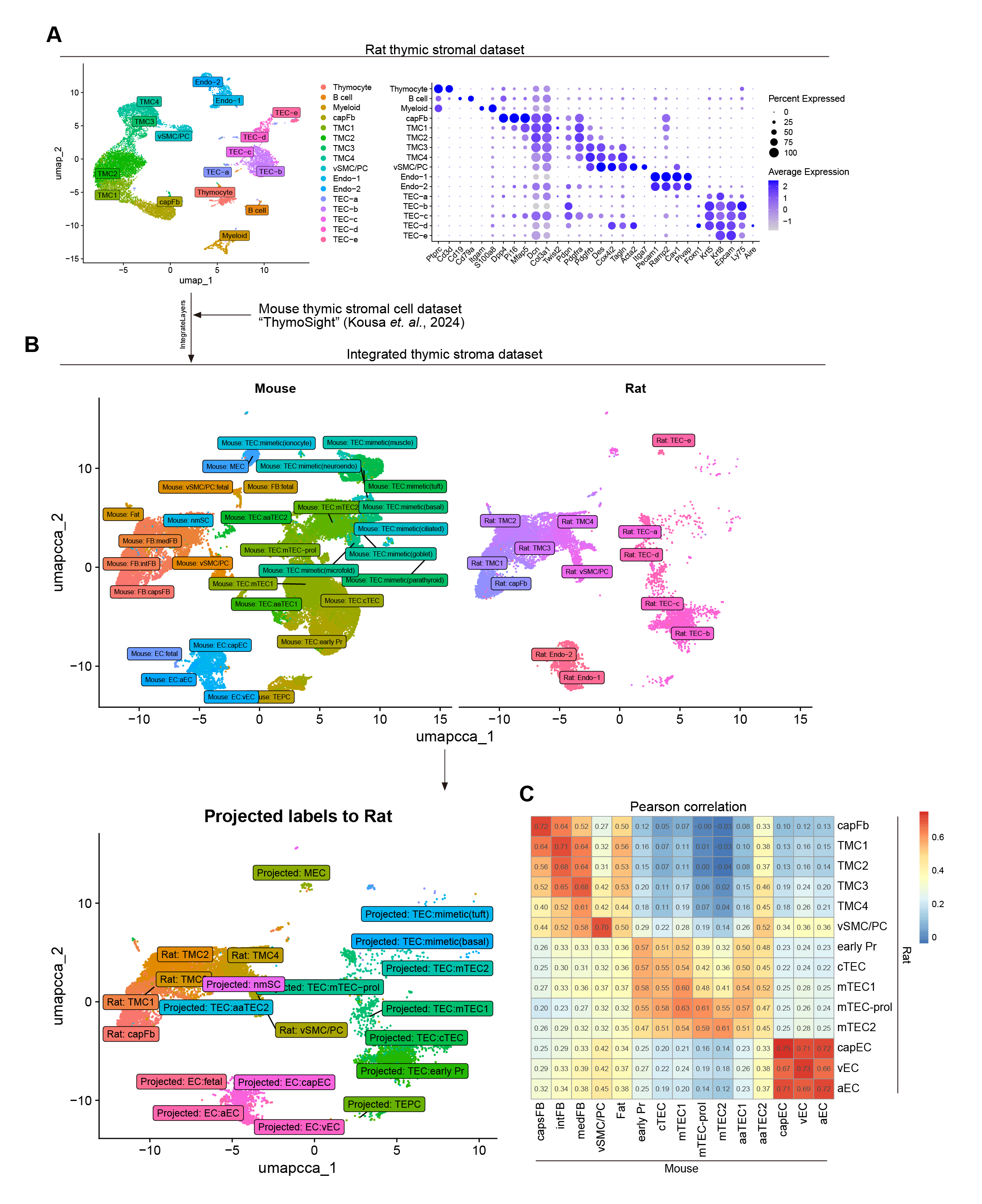

### FigS3.png

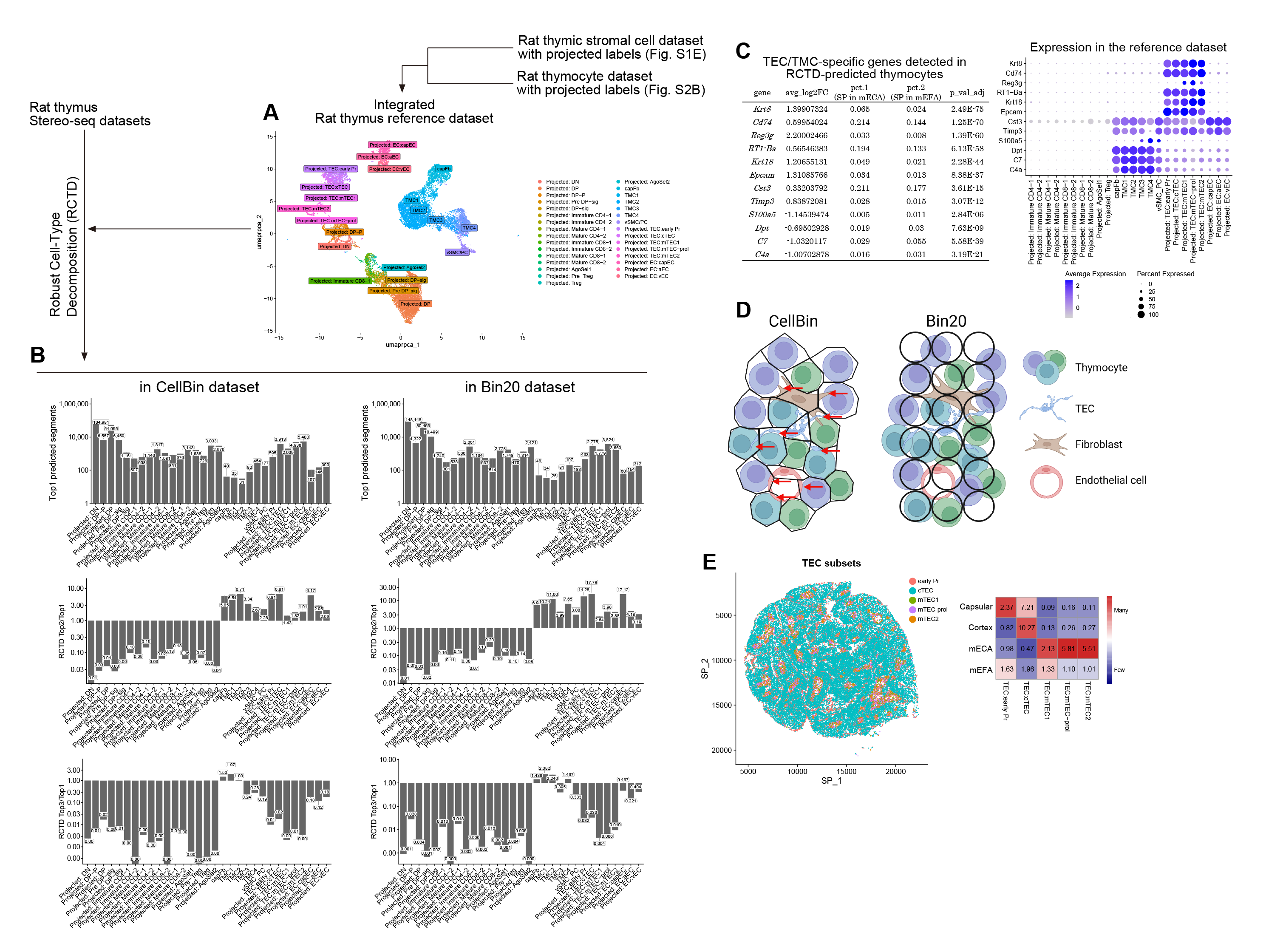

### FigS4.png

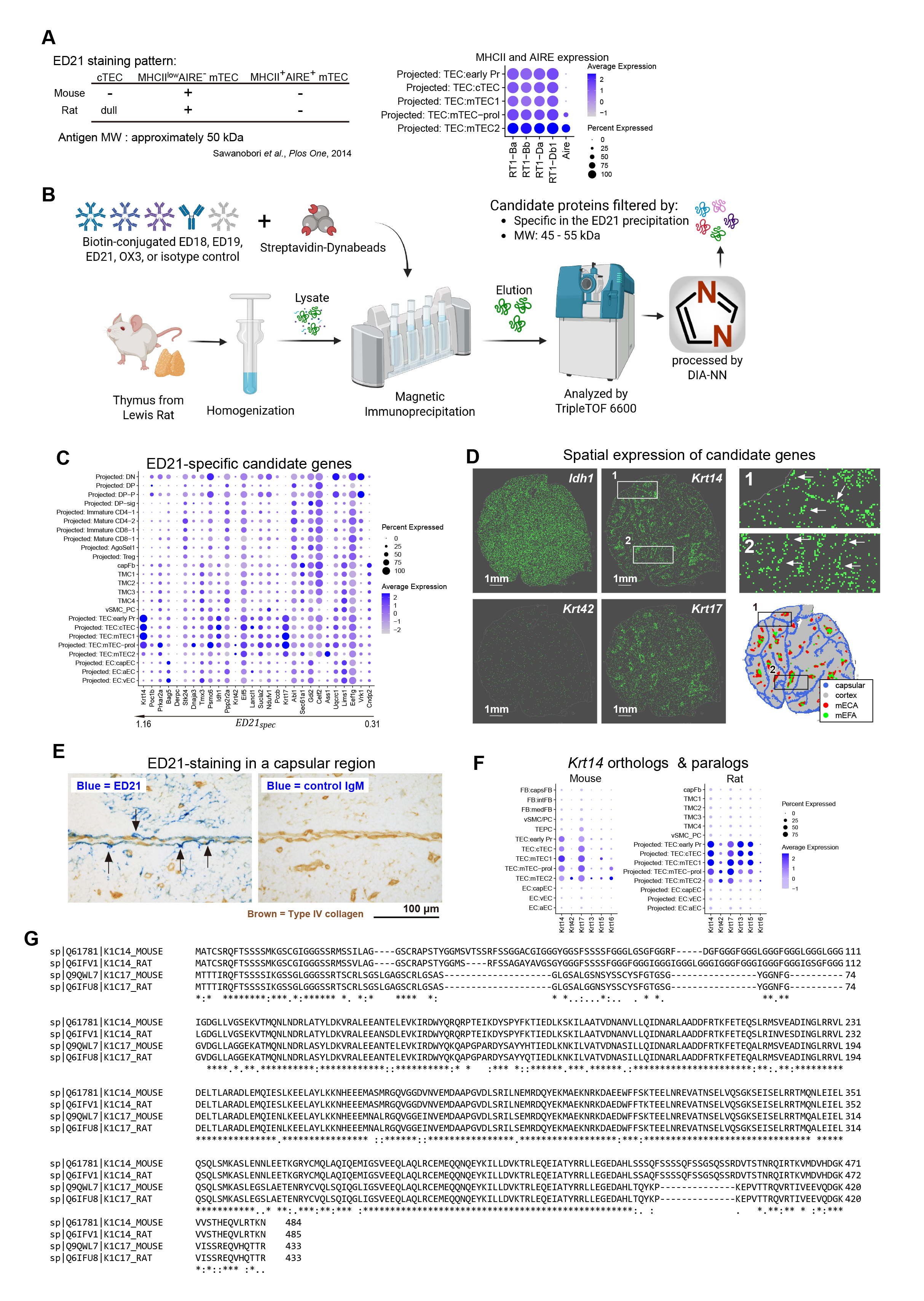

### mEFA-spatial analysis paper graphic abstract.png

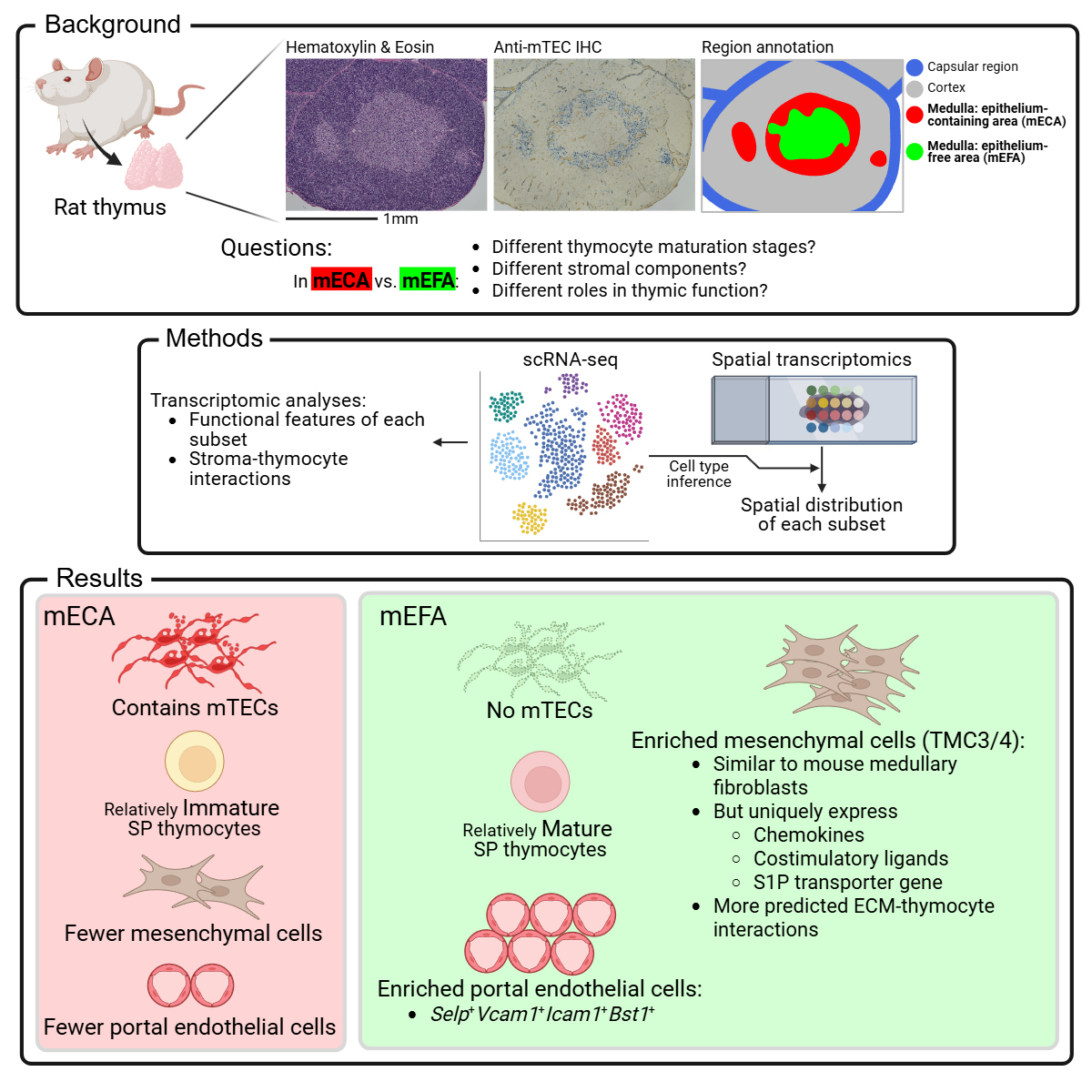
